# Antibiotic-induced shifts in fecal microbiota density and composition during hematopoietic stem cell transplantation

**DOI:** 10.1101/606533

**Authors:** Sejal Morjaria, Jonas Schluter, Bradford P. Taylor, Eric R. Littmann, Rebecca A. Carter, Emily Fontana, Jonathan U. Peled, Marcel R.M. van den Brink, Joao B. Xavier, Ying Taur

## Abstract

**Background:** Dramatic microbiota changes and loss of commensal anaerobic bacteria are associated with adverse outcomes in hematopoietic cell transplantation (HCT) recipients. In this study, we demonstrate these dynamic changes at high-resolution through daily stool sampling and assess the impact of individual antibiotics on those changes.

**Methods:** We collected 272 longitudinal stool samples (with mostly daily frequency) from 18 patients undergoing HCT and determined their composition by multi-parallel 16S rRNA gene sequencing, as well as density of bacteria in stool by qPCR. We calculated microbiota volatility to quantify rapid shifts and developed a new dynamic systems inference method to assess the specific impact of antibiotics.

**Results:** The greatest shifts in microbiota composition occurred between stem cell infusion and reconstitution of healthy immune cells. Piperacillin-tazobactam caused the most severe declines among obligate anaerobes.

**Conclusions:** Our approach of daily sampling, bacterial density determination and dynamic systems modeling allowed us to infer the independent effects of specific antibiotics on the microbiota of HCT patients.

## Background

Patients with a range of hematologic malignancies can be treated and potentially cured by hematopoietic cell transplantation (HCT). Prior to HCT, chemotherapy and/or total body irradiation are performed to deplete the cancerous cells. These treatments, combined with simultaneous antibiotic administration, compromise immune defenses, damage the mucosal epithelium, and deplete the native intestinal microbiota, facilitating the emergence of antibiotic-resistant organisms and increasing the risk of infections [1, 2]. Microbiome studies using fecal samples collected from allogeneic (allo-) HCT patients have previously revealed that patients experience severe reduction in the relative abundance of commensal bacteria over the course of treatment. This loss can result in blooms of potentially pathogenic microbial species [3], leading to downstream complications such as infections and graft-versus-host disease [4–6]. In particular, the loss of obligate anaerobic commensal bacteria such as Clostridia and Bacteroidetes negatively influence HCT outcomes shown in both animal models and humans [4, 7, 8].

Yet, the relative degree and manner in which various antibiotics and conditioning regimens contribute to microbiota disruption is still not well-described. In previous studies examining the microbiota changes in HCT patients, stool samples were either collected approximately once per week or at a limited number of time points [3–5, 9, 10]. Though these studies helped to form the foundation of our current understanding of microbiota disruption during all-HCT, we posit that a more frequent stool sample collection scheme, combined with dynamic modeling, would be beneficial for providing a higher resolution view of microbiota compositional changes over time, and where individual antibiotic effects can be discerned. Additionally, stool samples from many previous studies were characterized only in terms of relative abundance using 16S sequencing which does not allow quantitative calculations of species loss [11] and therefore, could potentially hamper attempts to quantitatively assess the effects of antibiotics on the microbiota.

In this study, we collected near-daily stool samples from 18 HCT recipients, for which we analyzed total species abundance by combining 16S sequencing in conjunction with quantitative PCR (qPCR) of the 16S gene. We leveraged classical models of microbial growth with Bayesian regression techniques to quantify the impact of specific classes of antibiotics on anaerobic microbes representative of a ‘healthy’ gut. Importantly, our model can be extrapolated to clinically guide sequential drug treatments that minimize detrimental effects on commensal bacteria. Our results reveal how important commensal anaerobic microbial species are lost during HCT.

## Methods

### Study patients and fecal sample collection

We followed 18 adult patients undergoing auto-HCT or allo-HCT at Memorial Sloan Kettering Cancer Center (MSKCC) from July 2015 to January 2016. There were 7 female and 11 male patients; their ages range from 40 to 75. Fecal samples were collected longitudinally from each patient during their transplant hospitalization using a prospective institutional fecal biospecimen collection protocol (described previously [3]). For the majority of patients, daily collection began at the start of pre-transplant conditioning (7-10 days before hematopoietic cell infusion) and continued until discharge, typically a month after HCT. The study protocol was approved by the MSKCC institutional review board; informed consent was obtained from all subjects prior to sample collection.

### Transplantation Practices

At MSKCC, antimicrobial prophylaxis is given routinely to patients undergoing HCT. Subjects undergoing either auto- or allo-HCT are given oral (PO) ciprofloxacin two days prior to hematopoietic cell infusion, as prophylaxis against gram negative bacterial infections. Allo-HCT recipients are also given intravenous (IV) vancomycin as prophylaxis against viridans-group streptococci [12]. Antibiotic prophylaxis against *Pneumocystis jiroveci* pneumonia was generally administered using either trimethoprim-sulfamethoxazole, aerosolized pentamidine, or atovaquone; the time at which prophylaxis was initiated (during conditioning or after engraftment, defined as an absolute neutrophil count of ≥500 neutrophils/mm^3^ for three consecutive days) varied. In the event of a new fever during times of neutropenia, patients were usually started on empiric antibiotics, such as piperacillin-tazobactam, cefepime, or meropenem.

### Sample Analysis of Microbial Composition

Sample DNA was extracted and purified, and the V4-V5 region of the 16S rRNA gene was amplified with polymerase chain reaction using modified universal bacterial primers. Sequencing was performed using the Illumina Miseq platform [13] yielding paired-end reads with length up to 250bp. These reads were assembled, processed, filtered for quality, and grouped into operational taxonomic units of 97% similarity using the UPARSE pipeline [14]. Taxonomic assignment to species level was performed using nucleotide BLAST (Basic Local Alignment Search Tool) [15], with the National Center for Biotechnology Information RefSeq (refseq_rna) as the reference database [16]. We determined the copy number of 16S rRNA genes per gram of stool for each sample by quantitative polymerase chain reaction (qPCR) on total DNA extracted from fecal samples. We assessed microbial diversity using the inverse Simpson index (for additional experimental details and microbiome data availability, see Supplementary Methods; all data used in this study are available as an excel file).

### Analytic approach

We developed and employed a metric of ‘compositional volatility’ to quantify the rate of overall change in microbiota composition across adjacent samples in time. The volatility metric assesses overall community change by calculating the Manhattan distance between microbiota compositions, and ranges between 0 and 1. It can be interpreted as the fraction of community turnover when comparing pairs of consecutive samples within a single patient. The volatility, *V*, for the community grouped at a taxonomic level, I, was determined by the expression:

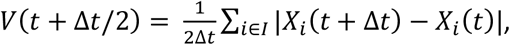

where Δt is the time in days between the consecutive samples and X_i_(t) is the relative abundance of taxa i at time t. We calculated volatility using relative abundances of microbes taxonomically grouped at the genus level. Theoretically, the most volatile points (V≈1), would correspond to complete microbiota replacements between two time-adjacent samples, whereby previously abundant genera would be completely replaced by different genera.

The total abundances of anaerobes were calculated by multiplying the summed relative abundances of obligate anaerobic taxa obtained by 16S sequencing (see extended methods for protocols and details) with the total copy numbers of 16S genes obtained via qPCR. To estimate the effects of different antibiotics on specific microbial groups, we calculated the log-difference of absolute anaerobe cell counts per gram of stool (wet weight) between two samples (deltas) which were at most two days apart and both within the first hospitalization. We used Bayesian regression techniques to parameterize a model of the logistic growth of the obligate anaerobe community, used similarly before [17]. Antibiotic effects on bacterial reproduction or death were modeled as independently modifying the anaerobe population growth rate. We also define two phases during HCT where loss of anaerobes might occur: Phase I = pre-HCT (i.e. 1 if a sample pair was obtained before stem cell infusion, 0 otherwise); Phase II = post-HCT but pre-engraftment (i.e. 1 if between day 0 and engraftment, 0 otherwise). We accounted for repeated samples from the same patient by including a random intercept term (1|P). Finally, we included a term that limits the otherwise exponential growth of anaerobes at high densities (the ‘capacity’ term of the logistic growth equation, with associated parameter βc). Changes in the anaerobe abundance, (N), were modeled as:

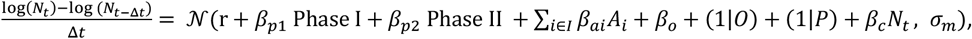

i.e. as a normally (𝒩) distributed variable that is a function of intrinsic growth rate (r), the effects of Phase I (β_p1_), Phase II (β_p2_) and the growth-rate changing effects, β_ai_, of empiric antibiotics (A_i_) and antibiotic prophylaxis (β_o_, *O*), with the residual uncaptured variance of the model, σ_m_). As for phase I and II, we constructed binary antibiotic covariate indicators (1 if an antibiotic was administered during the interval [t, t-Δt], 0 otherwise).

We considered a group effect of prophylactic antibiotics (β_o_) from which each individual prophylactic antibiotic (fluoroquinolones, vancomycin (IV), trimethoprim-sulfamethoxazole, atovaquone) could deviate (1|*O*, partial pooling of the effects of antibiotic prophylaxis). The empirical antibiotics piperacillin-tazobactam, meropenem, metronidazole, cephalosporins (gen.1-3), vancomycin (PO), cefepime and linezolid were considered without pooling.

We used uninformative priors (𝒩(0, 100^2^)) for the growth rate, empirical antibiotics, and the HCT treatment phases, and regularizing priors (𝒩(0, 10^−1^)) for the other parameters. This analysis produced posterior distributions for each parameter after “no U-turn” sampling 10,000 samples from 3 traces [18], each corresponding to an estimate of the degree of impact on obligate anaerobic bacterial populations.

We used the posterior parameter distributions to assess our model. We simulated the predicted changes for each patient’s timeline, starting with the first observed anaerobe count from that patient. We sampled 100 posterior predictions of anaerobe changes between timepoints, and used the mean predicted change for the calculation of the anaerobe count in the next timestep.

To describe the effect of realistic antibiotic treatment regimens on the group of commensal anaerobes in HCT patients, we compiled a list of all antibiotic administration courses as they occurred in our patient group, i.e. the duration of administration, the period when it was administered (e.g. Phase I or Phase II), and other co-administered antibiotics. We then repeatedly chose a random antibiotic course from this list, with replacement, and assigned parameters, chosen jointly from the posterior parameter value distributions, to our model. Then, starting from an initial, normalized density set to 1, we used the model to calculate the predicted fold-change of anaerobe density at the end of each antibiotic course. Aggregating all these fold-changes allowed us to calculate the average residual fraction of anaerobes after a ‘typical’ course of specific antibiotics.

## Results

### Description of study population and biospecimens

Our cohort consisted of 18 patients who underwent auto- or allo-HCT at MSKCC between July 2015 to January 2016. Clinical characteristics and stool microbiome data for each patient are shown in Figure 1. Patients underwent different types of HCT for a variety of conditions; 14 patients underwent allo-HCT and 4 underwent auto-HCT. The duration of transplant hospitalization ranged from 20 to 38 days, during which antibiotics were given for both prophylactic and treatment purposes. Throughout this period, we sought to collect fecal samples on a daily basis. A total of 272 samples were collected from the 18 patients (77%) of the total 352 total inpatient hospital days). Of those samples, 236 (87%) yielded 16S amplicons that could be sequenced.

**Figure 1:**
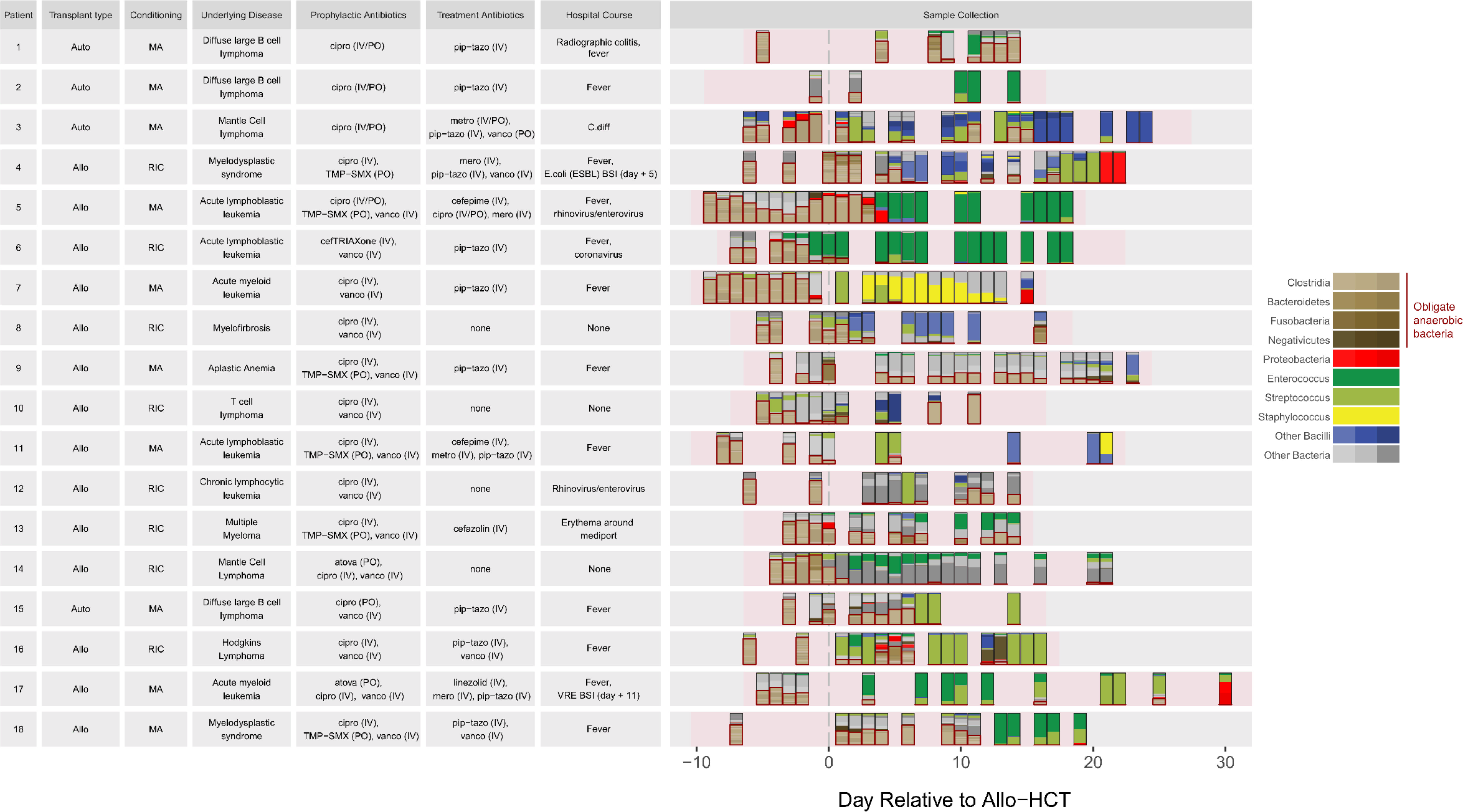
Clinical characteristics of all 18 HCT patients. Right panel depicts timing of stool collection during the course of transplantation (rectangles), relative to hematopoietic cell infusion (day 0), for each patient. Stacked colors represent each sample’s microbiota composition (based on 16S sequencing). Boxes are drawn around the anaerobes. Pink shading represents times of inpatient hospitalization. Abbreviations: Auto, autologous; Allo, allogeneic; RIC, reduced intensity conditioning; MA, myeloablative; IV, intravenous; PO, oral; cipro, ciprofloxacin; metro, metronidazole; pip-tazo, piperacillin-tazobactam; mero, meropenem; vanco, vancomycin; TMP-SMX, trimethoprim-sulfamethoxazole, C diff; *Clostridium difficile*; BSI; blood stream infection; VRE; vancomycin resistant enterococci

Consistent with previous findings [3, 5], all patients presented with high microbial diversity prior to HCT with species compositions that included diverse healthy anaerobic microbes; subsequent antibiotic administration caused large-scale changes to the intestinal microbiota with decreases specifically noted in intestinal diversity and bacterial population as a whole (Figures 1 and 2). This observation was noted in both all-HCT and auto-HCT patients (Supplementary Figure 1), coinciding with completion of pre-transplant conditioning and administration of broad-spectrum antibiotics around day 0 (Supplementary Figure 2). We focused on the obligate anaerobic bacterial species that fall in the classes, Clostridia/Negativicutes and the phyla, Fusobacteria/Bacteroidetes (Figure 3A) and noted a sharp decline in these bacterial populations post-HCT (Figure 3B).

**Figure 2:**
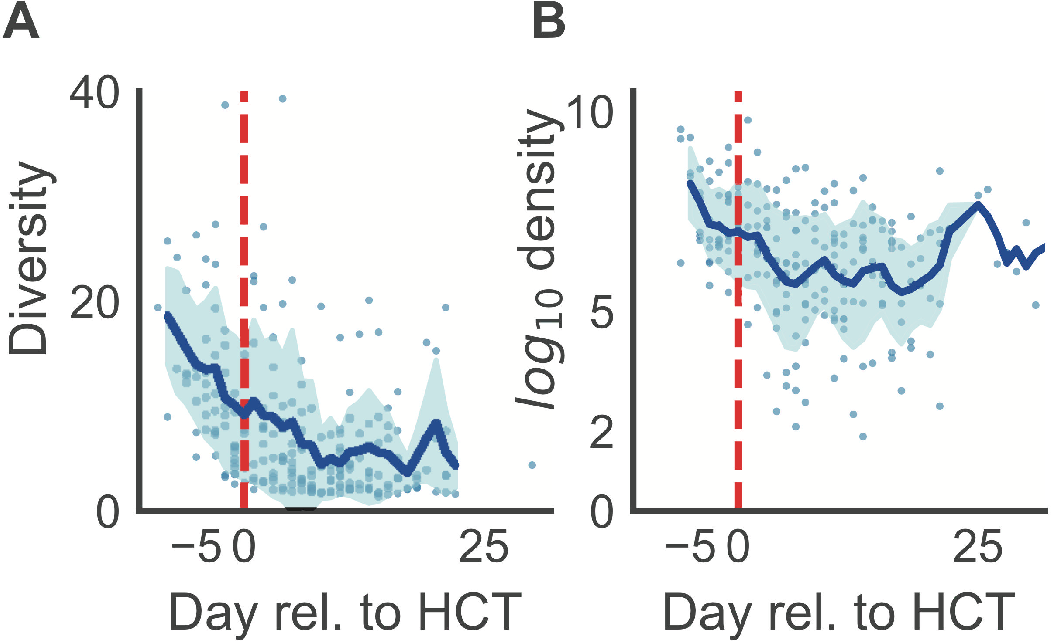
Microbiota changes in diversity and density during HCT. During conditioning, before hematopoietic cell transfusion (day 0, red vertical dashed line), the community diversity (A) of the microbiota in both allo- and auto-HCT patients declined rapidly. Similarly, the bacterial density declined, plotted as the total number of bacterial cells per gram of stool (B), and only mild recovery of cell counts was observed towards the latest days of hospitalization (and there, mostly in allo- patients, Supplementary Figure 1).

**Figure 3:**
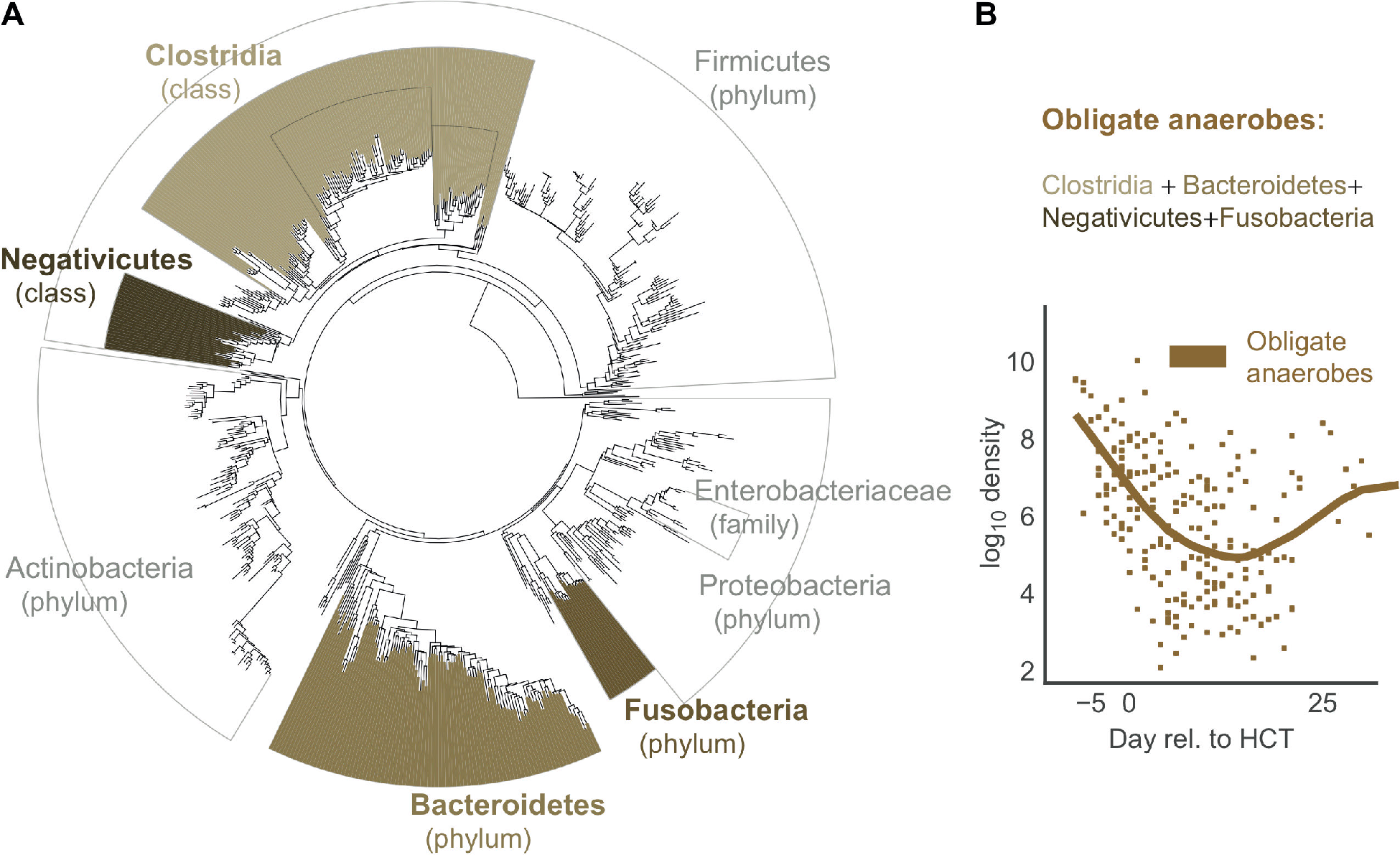
Timeline over the course of transplantation, for a single HCT patient (Patient 4). Antibiotic administration and chemotherapy regimen during conditioning (Phase I), post-HCT neutropenia before engraftment (Phase II), and post-engraftment (A). White blood cell counts across treatment with fever (thermometers) indicated (B). Relative abundances by 16S sequencing grouped at indicated taxon level during this patient’s HCT admission was collected almost daily (C). On day +3 the patient had a fever and received broad spectrum antibiotics as a result. Volatility quantifies the rate of change in microbiota composition across adjacent time points (D).

### Subject-level Microbiome changes

A single subject’s timeline (patient 4) is presented in Figure 4, showing medications and treatments, clinical data, and intestinal microbiome composition (Figure 4A-C). All other patients are shown in Supplementary Figure 3. We observed successive alterations of the intestinal microbiota in each patient over the course of transplantation. These dynamic changes seemed to correspond with specific changes in antibiotic administration. The loss of obligate anaerobic bacteria appeared to coincide more with the administration of certain types of antibiotics. Anaerobic microbes seemed relatively spared in some patients during periods where they remained only on prophylactic antibiotics (i.e. IV vancomycin, ciprofloxacin). In these 18 patients, we observed two patients with bloodstream infections, which were preceded by intestinal expansion of the corresponding pathogen (*Escherichia. coli* in Patient 5, and vancomycin-resistant Enterococcus faecium (VRE) in Patient 17; Supplementary Figure 3).

**Figure 4:**
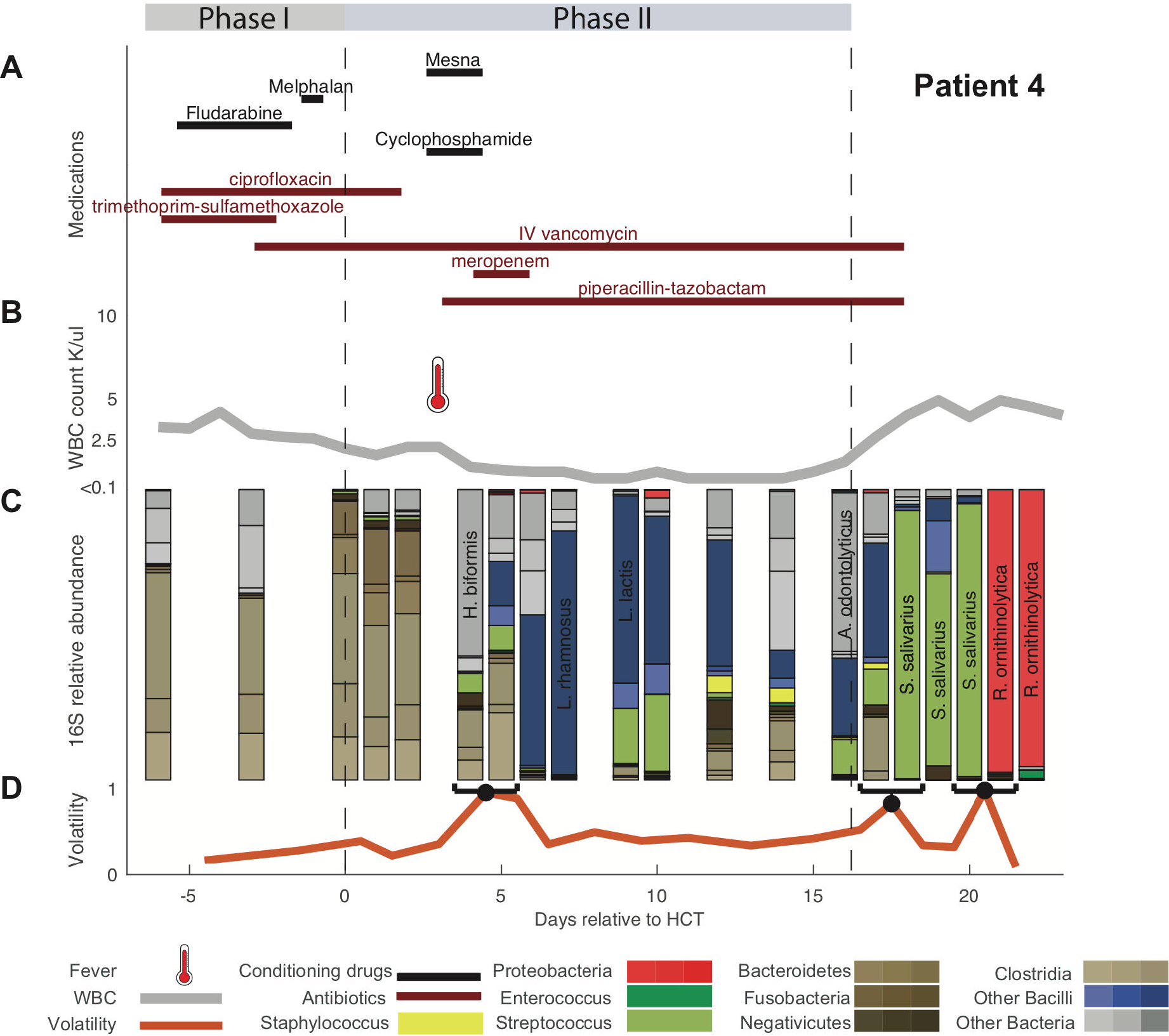
Obligate anaerobe grouping: Bacterial phylogenetic tree indicating in brown the groups we classified as commensal anaerobes (A). Average commensal anaerobe density across all patients (B, line: mean values per day using locally weighted scatterplot smoothing [31]).

To quantify day-to-day community shifts, we assessed the compositional volatility of the microbiota between daily intervals, reflecting the overall degree of compositional change over time (Figure 4D, Supplementary Figure 3). In our patients, microbiota volatility was on average highest immediately following transplant (Supplementary Figure 4).

### Antibiotic-induced loss of obligate anaerobic bacteria

Our model of microbial growth of obligate anaerobic bacteria identified piperacillin-tazobactam and meropenem as independently having the most detrimental impact on obligate anaerobes. Additionally, our model indicated a potential negative effect of metronidazole, cephalosporins (generation 1-3), and oral vancomycin, whereas fluoroquinolones, vancomycin (IV), and trimethoprim-sulfamethoxazole had no impact (Figure 5A). Furthermore, our model identified loss of obligate anaerobic bacteria when a patient experienced post-HCT neutropenia before neutrophil engraftment (Phase II), in addition to the effects of antibiotics. The model did not infer an exponential-growth limiting effect of the obligate anaerobe community onto itself (capacity). We were able to qualitatively capture the anaerobe dynamics of each patient’s time course, including major inflection points, by simulating each patient forwards in time starting with their first observed density of commensal anaerobes (Supplementary Figure 5).

**Figure 5:**
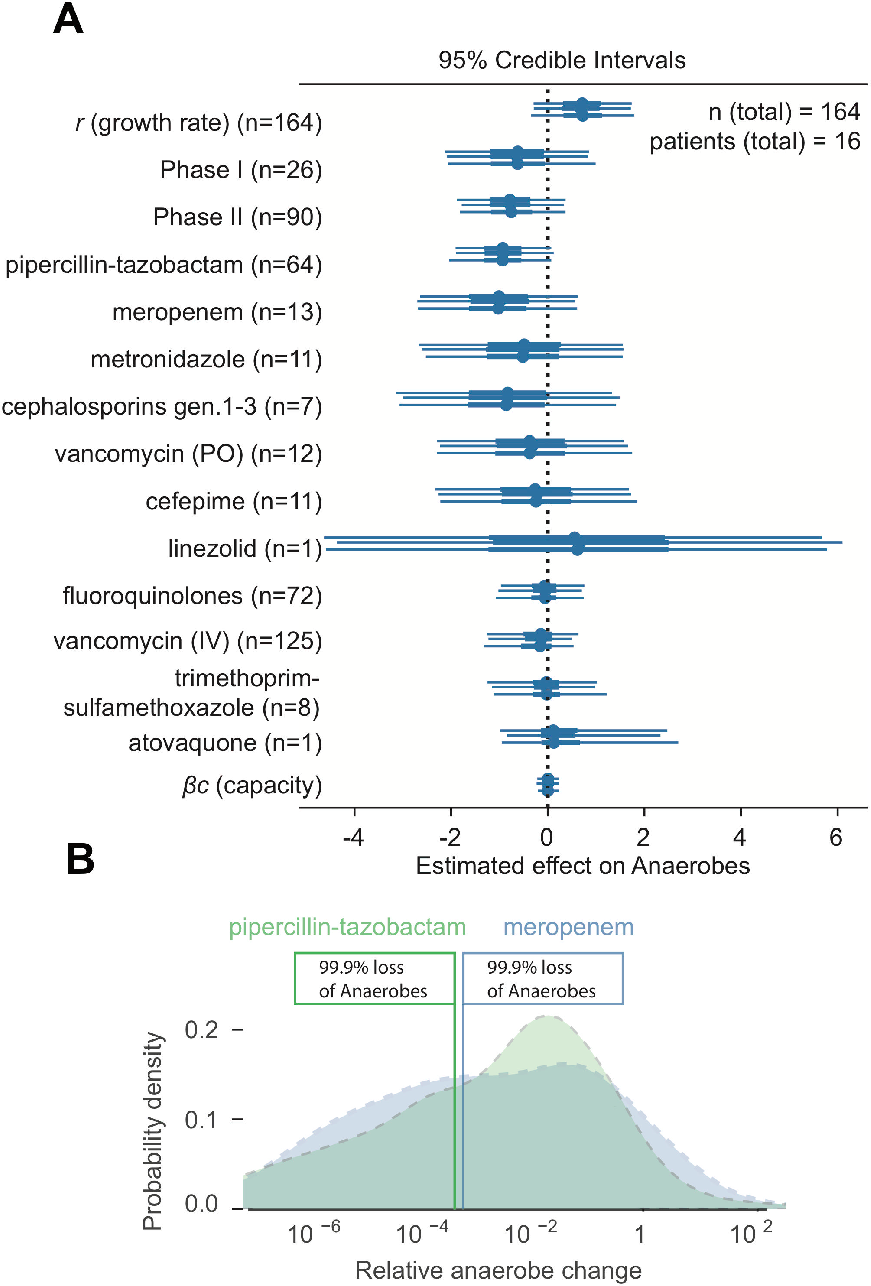
Specific antibiotic effects in HCT patients. Posterior parameter estimates from Bayesian linear regression of our model of antibiotic effects on obligate anaerobes. 95% Credibility intervals from three independent Markov Chain Monte Carlo traces with No-U-turn sampling. Distributions of predicted loss of anaerobes due to piperacillin-tazobactam and meropenem courses typical for our patient cohort (see methods for details, B).

We then predicted the effect of entire courses of piperacillin-tazobactam and meropenem as they occurred among our patients, i.e. including the effects of the period they were administered in (Phase I, Phase II or thereafter) as well as co-administered other antibiotics (Figure 5B, see methods). Due to differences in the duration of administrations and variation in co-administered antibiotics during these courses, the predicted loss of anaerobes was also variable (e.g. a longer course would yield a larger total loss of anaerobes). Yet, importantly, our model estimated that among our patients, courses with meropenem, and piperacillin-tazobactam each lead to >99% loss of obligate anaerobic bacteria (Figure 5B).

## Discussion

Under normal circumstances, the human intestinal microbiota is relatively stable over time, and largely consists of obligate anaerobic bacteria [19]. In contrast to this norm, our study shows dramatic day-to-day shifts in bacterial density and microbiota composition for HCT patients undergoing antibiotic treatment. Given the frequent administration of broad-spectrum antibiotics and the impact of HCT on immune defenses, mucosal epithelial integrity and dietary intake, these extreme shifts in microbiota-composition are perhaps unsurprising. In this study, our goal was to contextualize these compositional microbiota changes in terms of response to specific antibiotics [20]. We believe that a precise understanding and appreciation of antimicrobial influences on the microbiota can help inform antibiotic decision-making within the clinical setting, in order to protect against microbiota disruption and pathogen invasion. Our study is a first step towards the goal of minimizing unintended collateral damage to the commensal microbiota through the improved use of antibiotics.

Here, we show that patients can differ in terms of the rapidity, magnitude and quality of microbiota alterations, which likely reflect differences in baseline microbiota composition, exposure to antibiotics, and degrees of immune compromise and epithelial damage. The frequent stool sampling coupled with 16S qPCR and detailed clinical data, allowed us to now quantify the impact of specific antibiotics on microbiota composition. In our modeling approach, we incorporated absolute measures of 16S abundance and therefore were able to study microbiome dynamics as rate changes. This would be impossible with 16S relative abundance, and the resulting covariance bias could make it impossible to disentangle loss of anaerobes from increases in other taxa [21]. In some instances, our goal of daily stool collection was not well-met, leading to interval censoring.

We decided to analyze obligate anaerobic bacteria together in a single group (Clostridia, Bacteroidetes, Negativicutes and Fusobacteria) given their ties to healthy immunity and colonization resistance [4, 8, 10]. The premise that obligate anaerobic bacteria represented the vast majority of the normal colonic flora and conferred colonization resistance is rooted in pre-microbiome observations using culture-based methods, some as far back as 50 years ago [22]. Several studies have showcased their critical role in maintaining intestinal immune homeostasis [23–25] and association with good clinical outcomes [26, 27]. Admittedly, it is not known to what degree being ‘anaerobic’ approximates a truly beneficial microbiota. That said, by attempting to combine and study obligate anaerobic bacteria as we did, we were able to better align our results with existing clinical knowledge [28]; our model identified piperacillin-tazobactam and meropenem as major causes of obligate anaerobe loss, while metronidazole showed a propensity for obligate anaerobe killing, but to a lesser effect. A more robust killing effect on anaerobes by metronidazole may not have been seen because too few patients got this antibiotic (2), and it was often given during phase I and phase II, which may have offset its effect.

Our approach also demonstrated a degree of anaerobic impact from cephalosporins (generation 1-3). Although not traditionally thought to treat anaerobic infections, ceftriaxone and cefazolin have shown to have some activity against *Clostridium* spp. [29]. The killing potential of cephalosporins could also be explained by confounding factors such as its concurrent administration with specific conditioning treatments during Phase I that were not included explicitly in our model (patients 6 and 13; Supplementary Figure 3).

Indeed, we observed significant anaerobic loss during the time window of Phase II, i.e. post-HCT but prior to neutrophil engraftment day. Independent of antibiotic effects, these results may reflect direct conditioning-related effects that directly impact intestinal homeostasis, which we suspect occurs to at least to a certain degree. These effects could consist of damage from chemotherapy and/or radiation, either to the intestinal mucosa (thereby impacting niche factors and microbiome homeostasis), or to anaerobic bacteria directly within the lumen.

HCT exposes the intestinal microbiota to a wide variety of environmental changes including a new residence and diet and creates a complex ecosystem that is difficult to model. However, despite this being a pilot study of HCT patients, we feel our model performed well, and provided promising results largely consistent with our clinical impressions. Discerning the individual effects of different antibiotics and chemotherapy on the microbiota was a challenge, as simultaneous drug administrations are common, but we are confident with our model estimations for individual antibiotic effects. Still, our complex patient population and small sample size meant some of our results consisted of wide credibility intervals, which estimate population parameters with lower precision. Our model therefore predicted the time courses of patients qualitatively, capturing inflection points of major anaerobe loss rather than predicting time courses with high quantitative accuracy. To bypass these potential limitations, we will continue to accumulate high frequency, quantitative microbiome data in conjunction with clinical metadata to better predict the individual effects of each drug.

Understanding collateral damages to the microbiota is not only relevant to prevent microbiome dysbiosis-related disease, but also to prevent the rise of antibiotic resistant pathogens [20, 30]. A better understanding of the dynamics that render a complex microbiota permissive to pathogen expansion has the potential to shape and improve basic principles of antibiotic stewardship.

## Footnotes

### Meeting(s) where the information has previously been presented

The information contained in this paper has been presented at previous National meetings.

### Corresponding author contact information

Ying Taur, MD, MPH, Correspondence: Ying Taur, MD, MPH, Memorial Sloan-Kettering Cancer Center, 1275 York Avenue, Box 9, New York, NY 10065

## Supplementary Figures and Captions

**Supplementary Figure 1:**
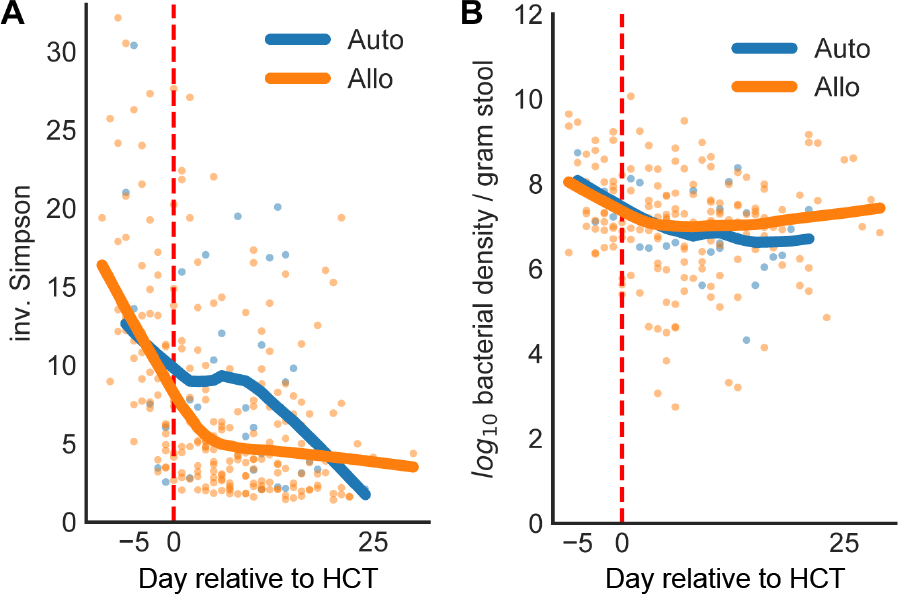
Microbiota destruction during HCT. During conditioning, before stem cell transfusion (day 0), the community diversity (A) of the microbiota in both allo- and auto-HCT patients tended to decline rapidly. Similarly, the bacterial density, measured as the total number of bacterial cells per gram of stool (B), also declined, with slight recovery of cell counts in allo-HCT patients during later days of hospitalization.

**Supplementary Figure 2:**
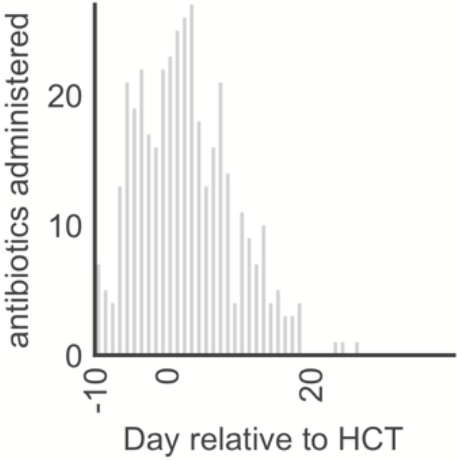
Total counts of antibiotics prescribed per day relative to HCT, summed over all 18 patients.

**Supplementary Figure 3:** Timelines of all patients (see Figure 4). Available as combined supplementary file “supplementary_figure3.pdf”.

**Supplementary Figure 4:**
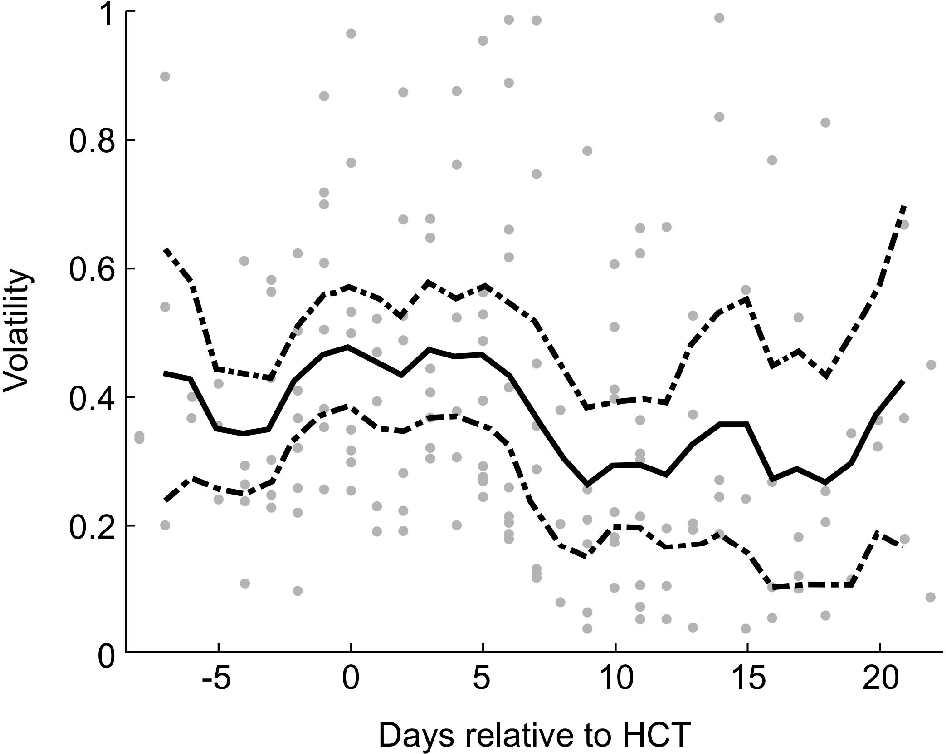
Microbiota volatility per day relative to HCT between daily samples. The line shows a three day rolling average, dashed lines indicate the 95% confidence intervals of the mean.

**Supplementary Figure 5:**
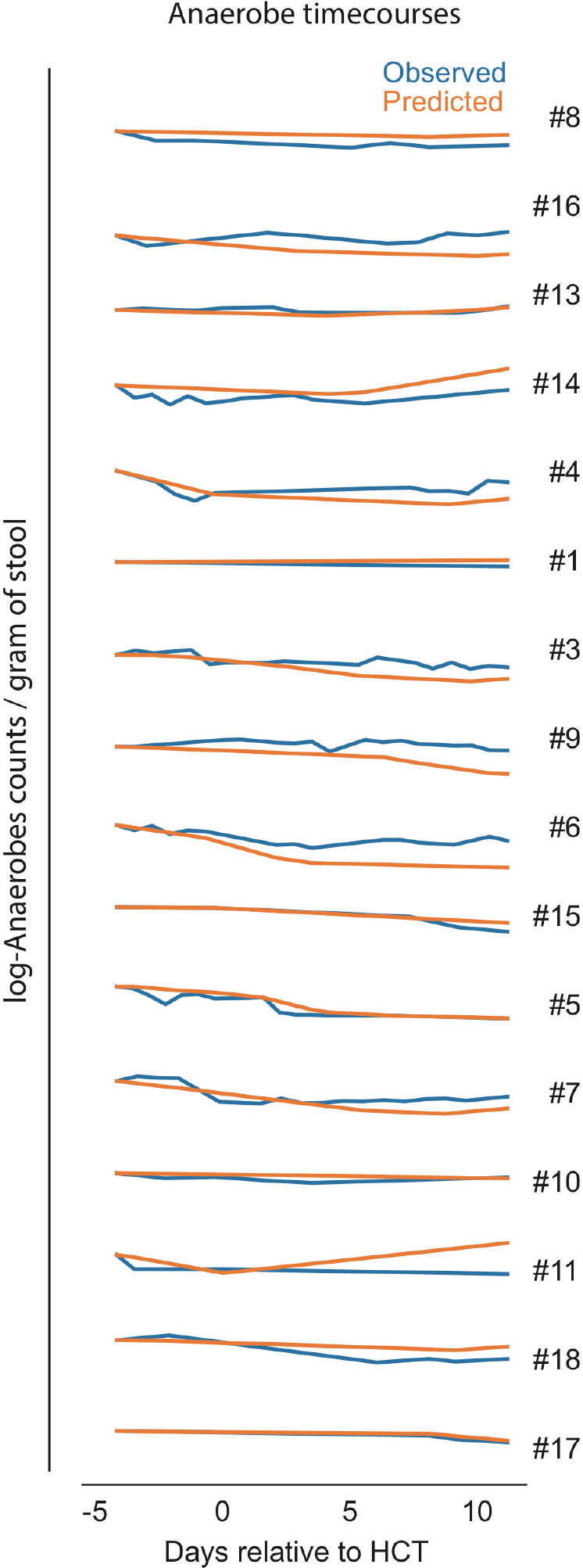
Model predictions. A) Histogram of observed (blue) fold changes in anaerobe counts followed a similar distribution to the posterior predictions (orange). B) Starting from each patient’s first observed anaerobe counts, we simulated forwards in time and plot the average predicted anaerobe time course (orange) against the observed (blue).

## Supplemental Methods

### Sample preparation and sequencing protocols

#### DNA extraction

Briefly, a frozen aliquot (≈100 mg) of each sample was suspended, while frozen, in a solution containing 500 μl of extraction buffer (200 mM Tris, pH 8.0/200 mM NaCl/20 mM EDTA), 200 μl of 20% SDS, 500 μl of phenol:chloroform:isoamyl alcohol (25:24:1), and 500 μl of 0.1-mm-diameter zirconia/silica beads (BioSpec Products). Microbial cells were lysed by mechanical disruption with a bead beater (BioSpec Products) for 2 min, after which two rounds of phenol:chloroform:isoamyl alcohol extraction were performed. DNA was precipitated with ethanol at −80 degrees and resuspended in 200 μl of TE buffer with 100 mg/ml RNase. The isolated DNA was subjected to additional purification with QIAamp mini spin columns (Qiagen).

#### 16S rDNA amplification and Illumina Sequencing

For each sample, duplicate 50-μl PCR reactions were performed, each containing 50 ng of purified DNA and a master mix of 0.2 mM dNTPs, 1.5 mM MgCl2, 2.5 U Platinum Taq DNA polymerase, 2.5 μl of 10X PCR buffer, and 0.5 μM of each primer designed to amplify the V4-V5: 563F (5’-nnnnnnnn-NNNNNNNNNNNN-AYTGGGYDTAAAGNG-3’) and 926R (5’- nnnnnnnn-NNNNNNNNNNNN-CCGTCAATTYHTTTRAGT-3’). A unique 12-base Golay barcode (Ns) precede the primers for sample identification [12] and 1-8 additional nucleotides were placed in front of the barcode to offset the sequencing of the primers. Cycling conditions were 94°C for 3 minutes, followed by 27 cycles of 94°C for 50 seconds, 51°C for 30 seconds, and 72°C for 1 minute. 72°C for 5 min is used for the final elongation step. Replicate PCR products were pooled and amplicons were purified using the Qiaquick PCR Purification Kit (Qiagen). PCR products were quantified and pooled at equimolar amounts before proceeding with library preparation following the Illumina TruSeq Sample Preparation protocol. The completed library was sequenced on an Illumina Miseq platform following the Illumina recommended procedures with a paired end 250 × 250 bp kit.

#### Sequence processing

Paired end reads were assembled, processed, and grouped into operational taxonomic units (OTUs) of 97% similarity using the UPARSE pipeline [13]. Sequences were error-filtered, using maximum expected error (Emax=1). Taxonomic assignment to species level was performed for representative sequences form each OUT; this was achieved by using a custom python script incorporating nucleotide BLAST (Basic Local Alignment Search Tool), with the National Center for Biotechnology Information RefSeq as the reference training set [14]. We obtained a total of 4,055,808 high-quality 16S rRNA gene-encoding sequences, with a mean of 12,887 sequences per sample. A phylogenetic tree was constructed by aligning representative sequences to SILVA 16S reference.

Sequence designations and identity scores were manually inspected for quality and consistency in terms of taxonomic structure and secondary matches. Based on our testing and comparisons using mock community data, we have found this approach to yield good robust species-level approximations for our candidate sequences. In particular, species-level classification of clostridial species such as Clostridium difficile improved greatly compared with other routine classification methods.

#### Total 16S quantification

Copy number of 16S rRNA genes for each sample was determined by quantitative PCR (qPCR) on total DNA extracted from fecal samples. Primers specific to the V4 - V5 region of the 16S gene 563F (5’- AYTGGGYDTAAAGNG-3’) and 926Rb (5’-CCGTCAATTYHTTTRAGT-3’) at 0.2 μM concentrations were used with the DyNAmo HS SYBR green qPCR kit (Thermo Fisher Scientific). In order to determine absolute abundances and copy numbers of the 16S gene of unknown samples, a standard was created by taking the V4 and V5 regions from *Escherichia coli* cloned into the *Invitrogen* TOPO pcr2.1 TA vector^AMP^. The plasmid and insert are 4318bp in length. Copies / μL of our standard is calculated and a total of 7 1:5 serial dilutions starting with 100,000,000 copies create the standard curves which we map our unknown samples against.

The cycling conditions were as follows: 95°C for 10 min, followed by 40 cycles of 95°C for 30 s, 52°C for 30 s, and 72°C for 30 s. 16s qPCR was performed on all stool samples in order to determine bacterial density in feces. We were unable to amplify bacterial 16S genes from 38 samples, suggesting that bacterial density in these samples was below the level of detection.

