## supplementary figure 3 for "Antibiotic-induced shifts in fecal microbiota density and composition during hematopoietic stem cell transplantation"

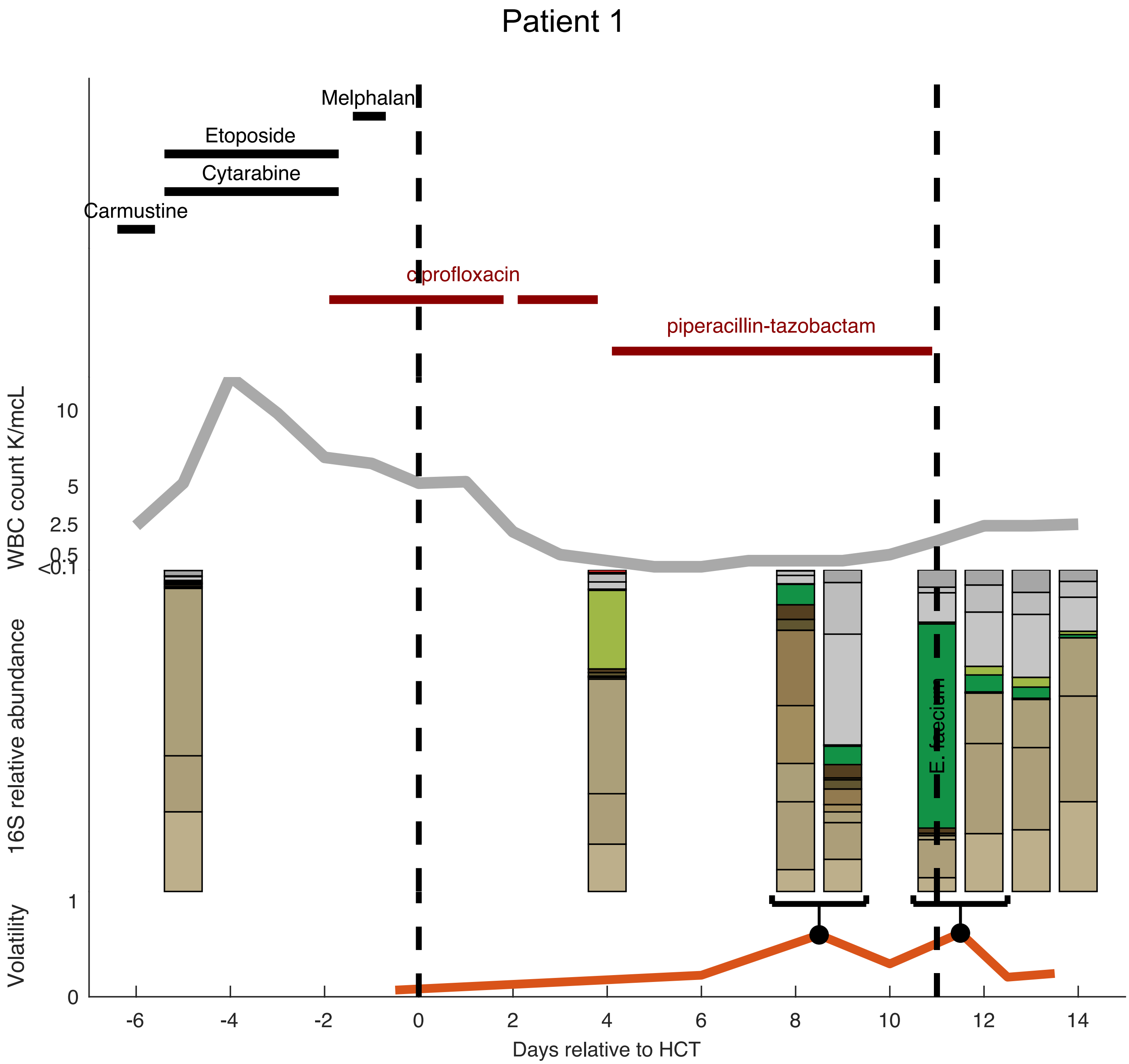

Patient 1
Melphalan
Etoposide
Cytarabine
Carmustine
ciprofloxacin
piperacillin-tazobactam
WBC count K/mcL
10
5
2.5
0.5
<0.1
16S relative abundance
E. faecium
Volatility
1
0
-6
-4
-2
0
2
4
6
8
10
12
14
Days relative to HCT

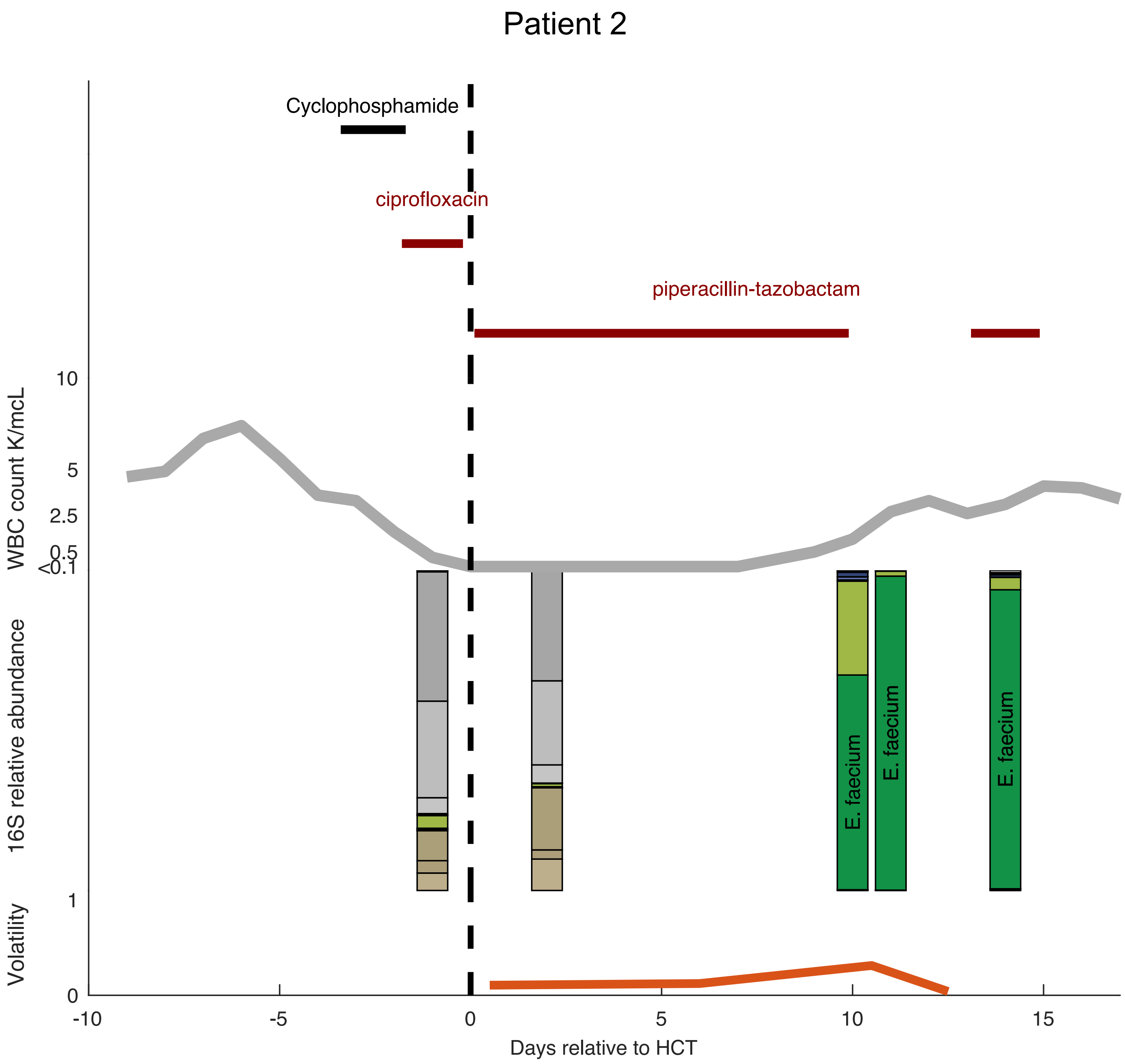

Patient 2
Cyclophosphamide
ciprofloxacin
piperacillin-tazobactam
WBC count K/mcL
10
5
2.5
0.5
<0.1
16S relative abundance
E. faecium
E. faecium
E. faecium
Volatility
1
0
-10
-5
0
5
10
15
Days relative to HCT

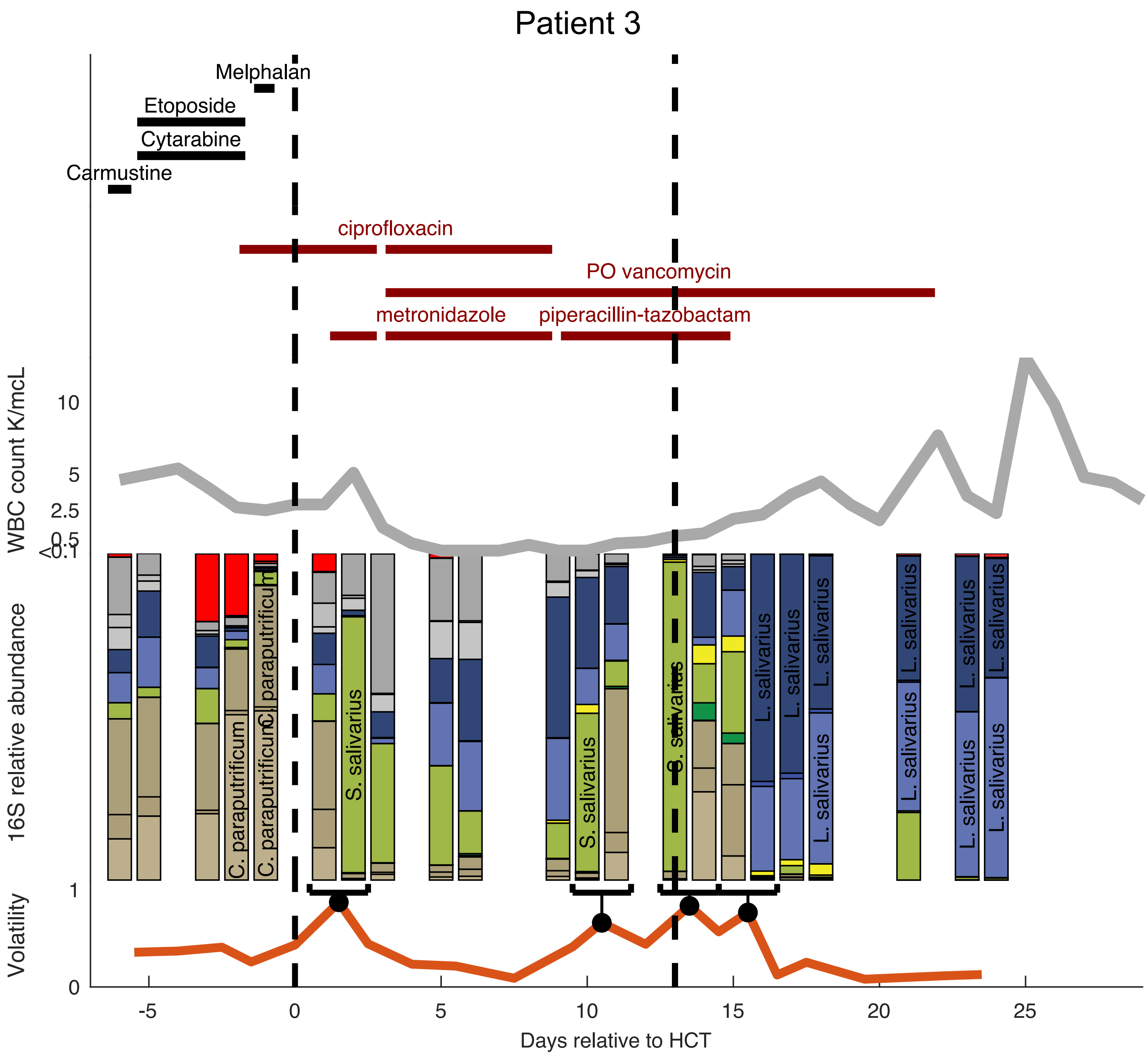

Patient 3
Melphalan
Etoposide
Cytarabine
Carmustine
ciprofloxacin
PO vancomycin
metronidazole
piperacillin-tazobactam
WBC count K/mcL
10
5
2.5
0.5
<0.1
16S relative abundance
C. paraputrificum
S. salivarius
L. salivarius
Volatility
1
0
-5
0
5
10
15
20
25
Days relative to HCT

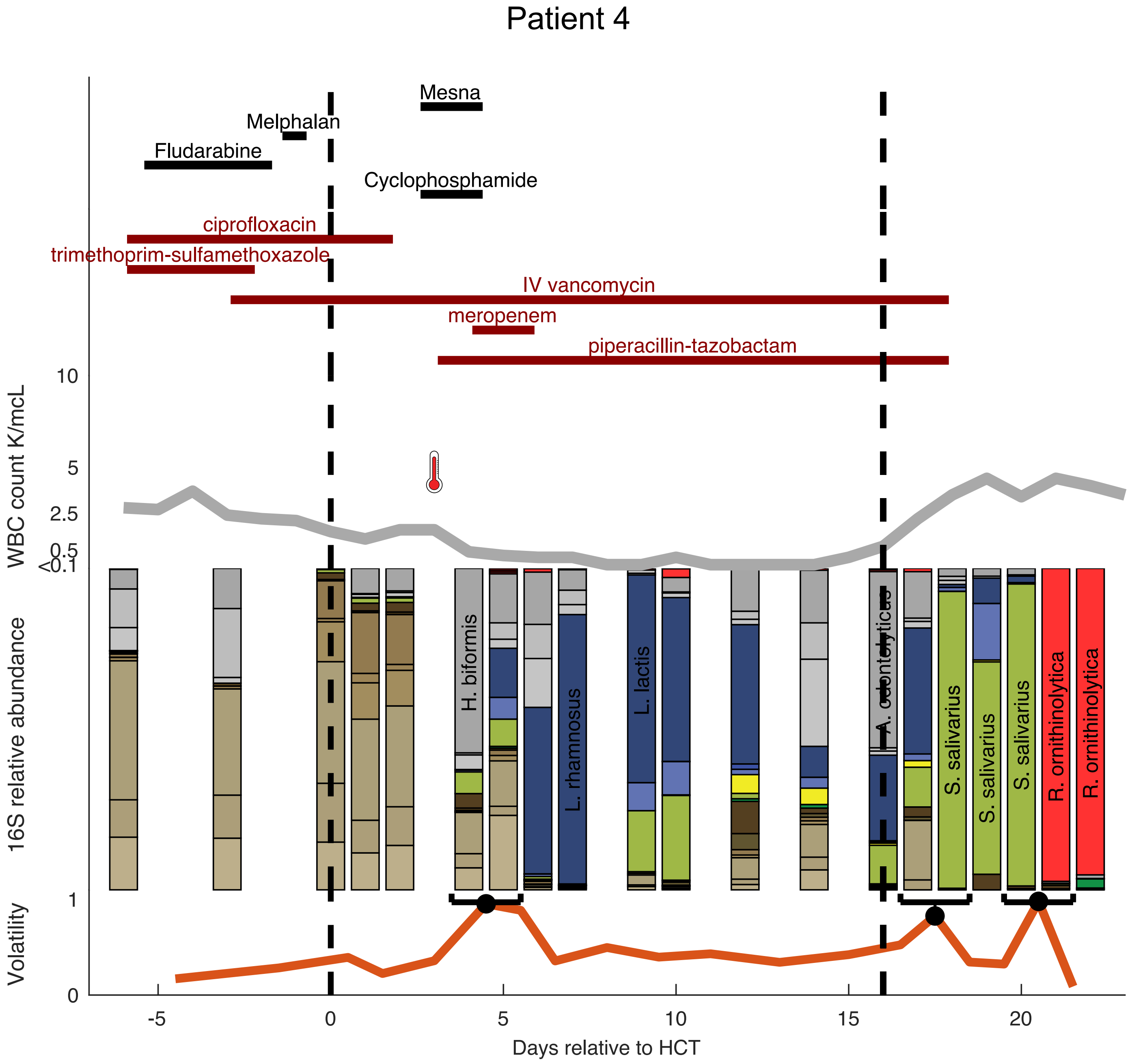
Patient 4
Mesna
Melphalan
Fludarabine
Cyclophosphamide
ciprofloxacin
trimethoprim-sulfamethoxazole
IV vancomycin
meropenem
piperacillin-tazobactam
WBC count K/mcL
10
5
2.5
0.5
<0.1
16S relative abundance
H. biformis
L. rhamnosus
L. lactis
A. odontolyticus
S. salivarius
S. salivarius
S. salivarius
R. ornithinolytica
R. ornithinolytica
Volatility
1
0
-5
0
5
10
15
20
Days relative to HCT

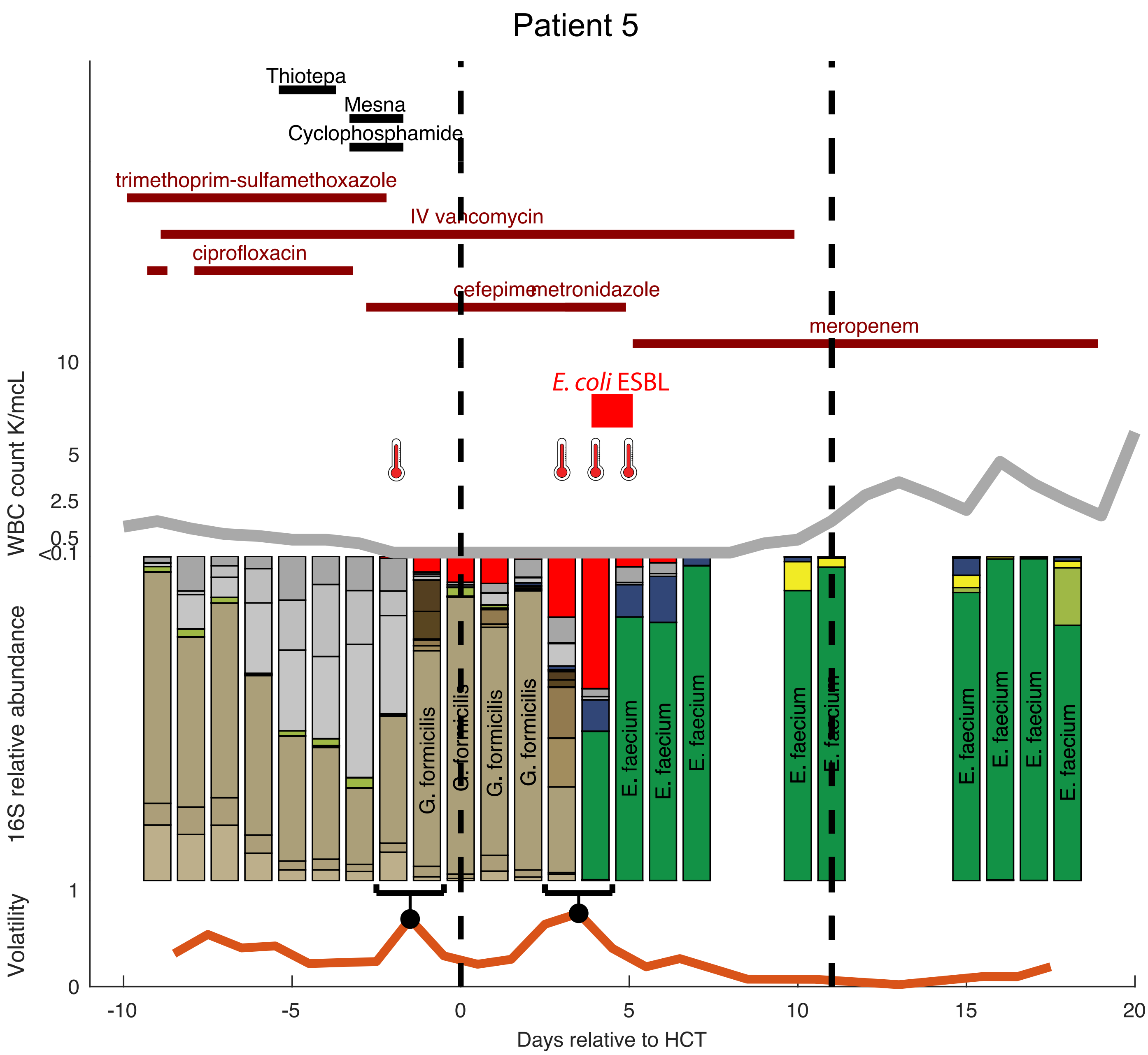
Patient 5
Thiotepa
Mesna
Cyclophosphamide
trimethoprim-sulfamethoxazole
IV vancomycin
ciprofloxacin
cefepime
metronidazole
meropenem
E. coli ESBL
WBC count K/mcL
10
5
2.5
0.5
<0.1
16S relative abundance
G. formicilis
G. formicilis
G. formicilis
G. formicilis
E. faecium
E. faecium
E. faecium
E. faecium
E. faecium
E. faecium
E. faecium
E. faecium
E. faecium
Volatility
1
0
-10
-5
0
5
10
15
20
Days relative to HCT

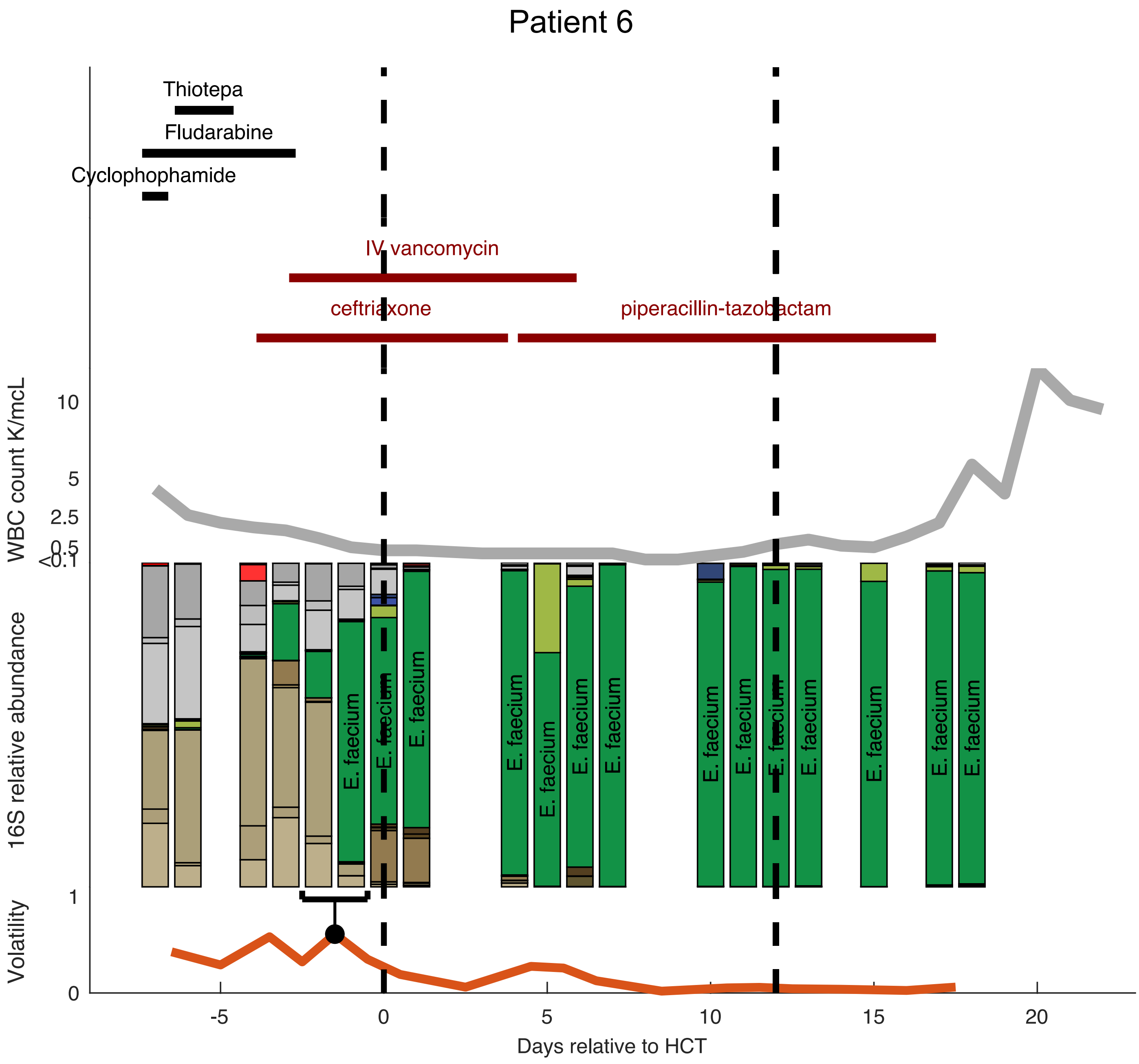

Patient 6
Thiotepa
Fludarabine
Cyclophophamide
IV vancomycin
ceftriaxone
piperacillin-tazobactam
WBC count K/mcL
10
5
2.5
0.5
<0.1
16S relative abundance
E. faecium
Volatility
1
0
-5
0
5
10
15
20
Days relative to HCT

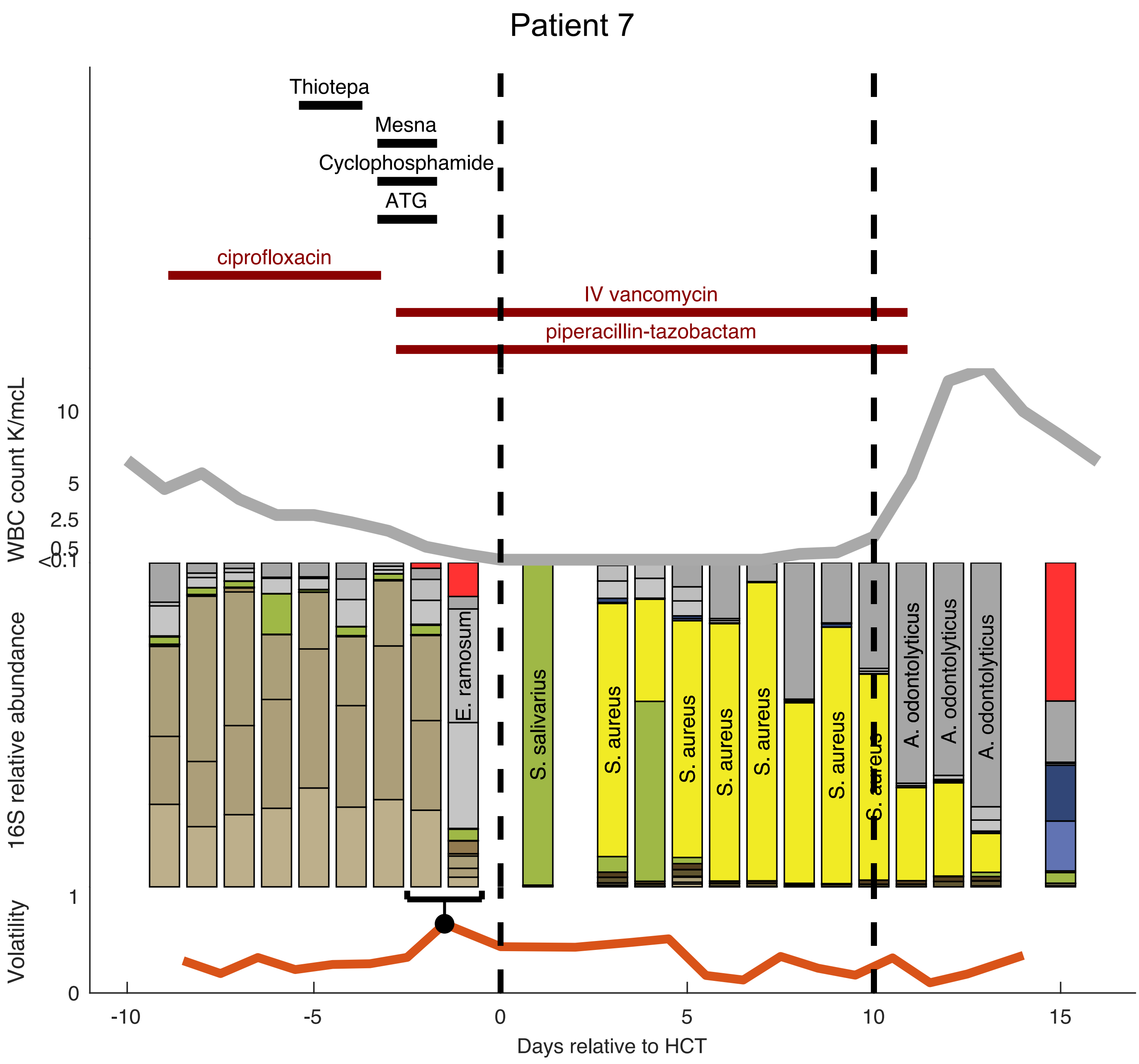

Patient 7
Thiotepa
Mesna
Cyclophosphamide
ATG
ciprofloxacin
IV vancomycin
piperacillin-tazobactam
WBC count K/mcL
10
5
2.5
0.5
<0.1
16S relative abundance
E. ramosum
S. salivarius
S. aureus
S. aureus
S. aureus
S. aureus
S. aureus
S. aureus
A. odontolyticus
A. odontolyticus
A. odontolyticus
Volatility
1
0
-10
-5
0
5
10
15
Days relative to HCT

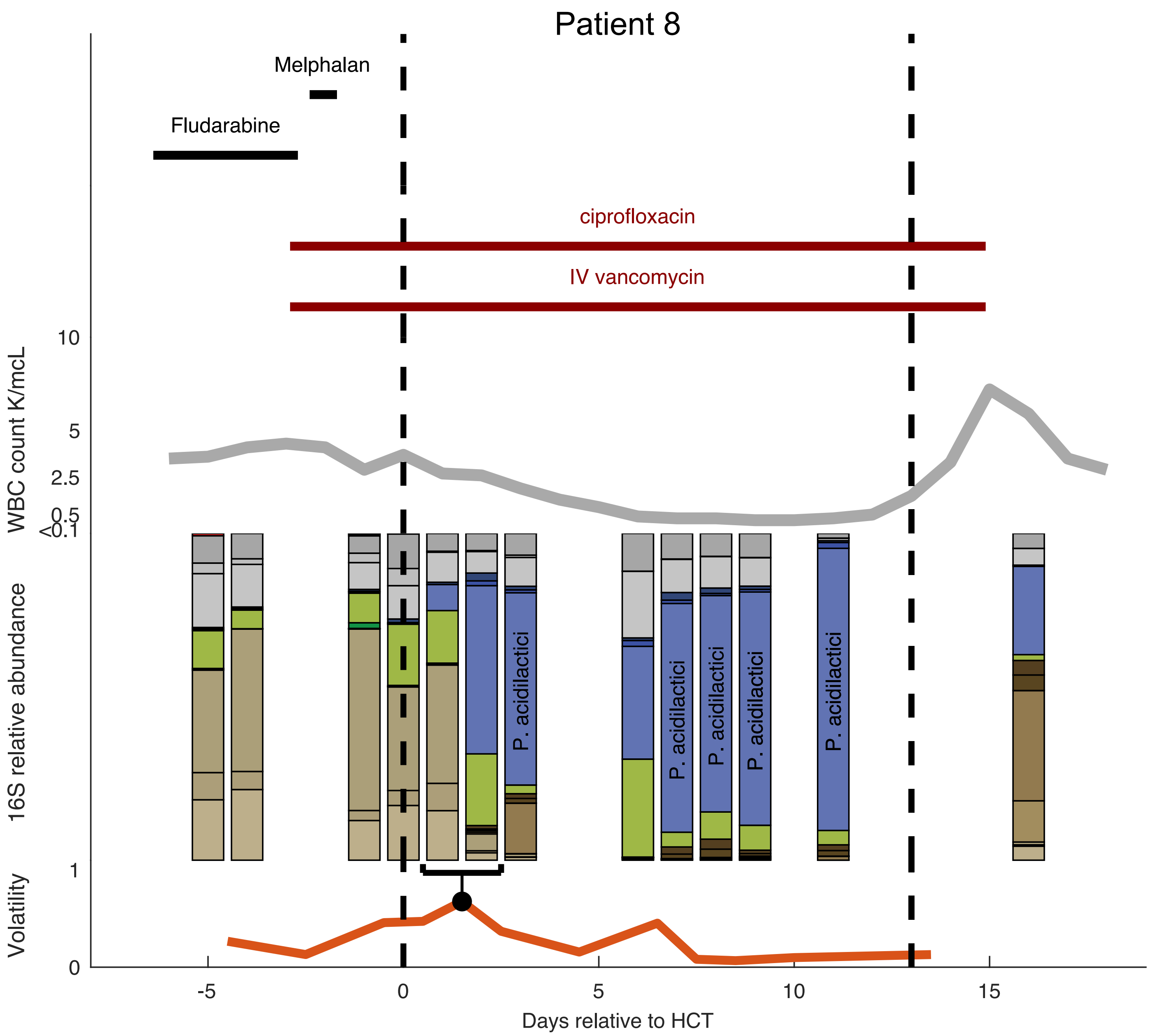
Patient 8
Melphalan
Fludarabine
ciprofloxacin
IV vancomycin
10
5
2.5
0.5
<0.1
WBC count K/mcL
16S relative abundance
P. acidilactici
P. acidilactici
P. acidilactici
P. acidilactici
P. acidilactici
1
0
Volatility
-5
0
5
10
15
Days relative to HCT

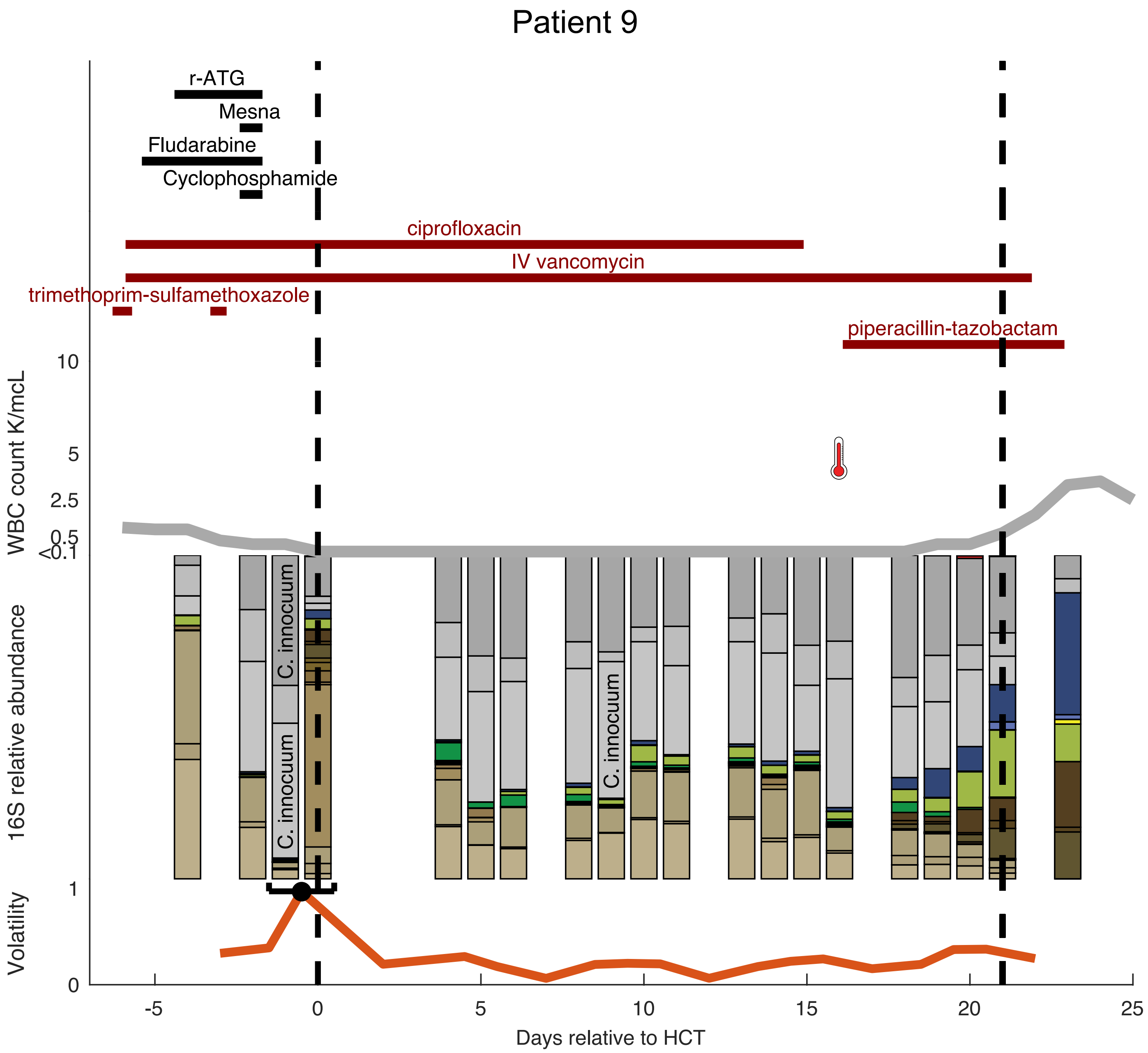
Patient 9
r-ATG
Mesna
Fludarabine
Cyclophosphamide
ciprofloxacin
IV vancomycin
trimethoprim-sulfamethoxazole
piperacillin-tazobactam
WBC count K/mcL
10
5
2.5
0.5
<0.1
16S relative abundance
C. innocuum
C. innocuum
C. innocuum
Volatility
1
0
-5
0
5
10
15
20
25
Days relative to HCT

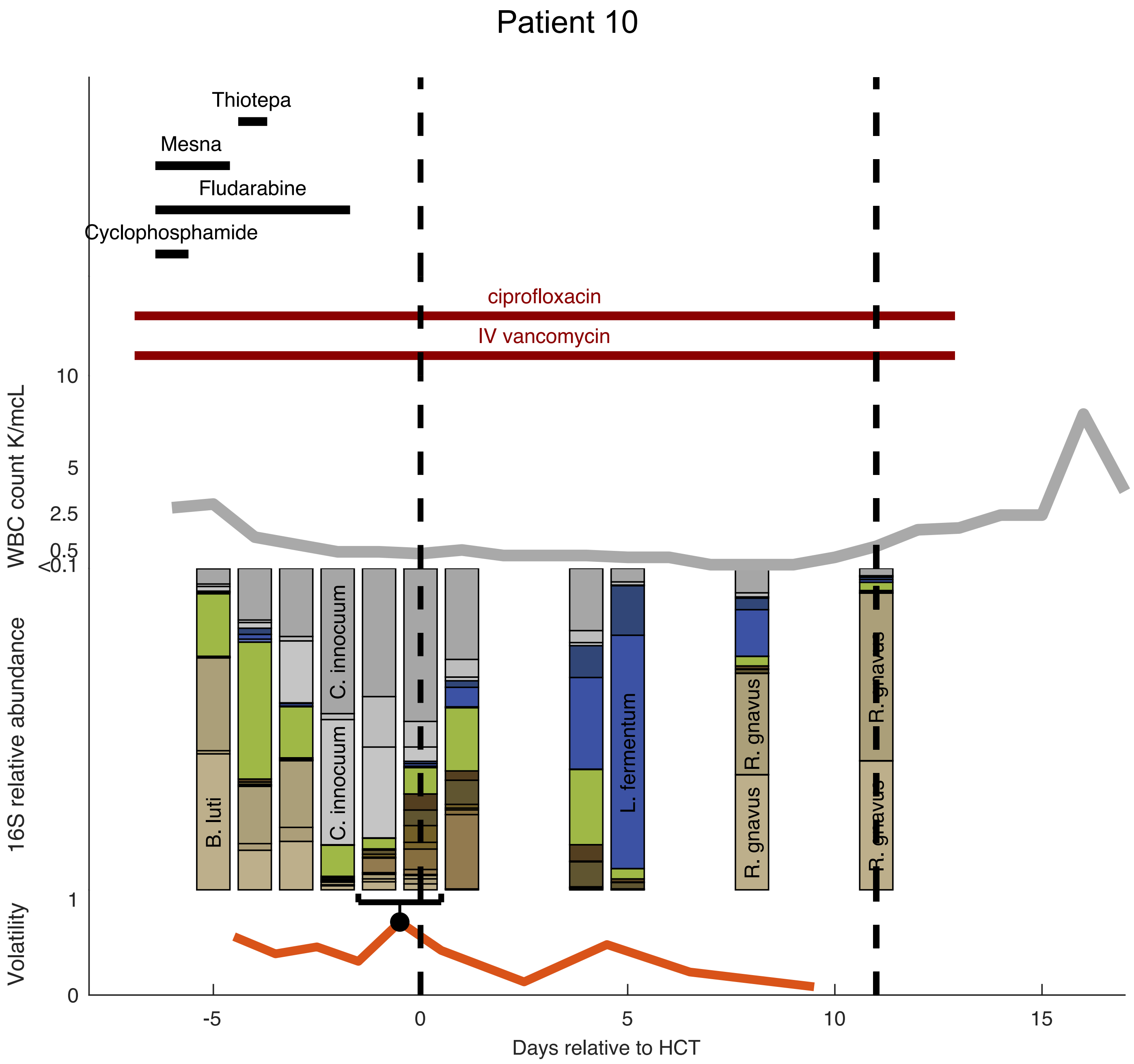

Patient 10
Thiotepa
Mesna
Fludarabine
Cyclophosphamide
ciprofloxacin
IV vancomycin
WBC count K/mcL
10
5
2.5
0.5
<0.1
16S relative abundance
B. luti
C. innocuum
C. innocuum
L. fermentum
R. gnavus
R. gnavus
R. gnavus
R. gnavus
Volatility
1
0
-5
0
5
10
15
Days relative to HCT

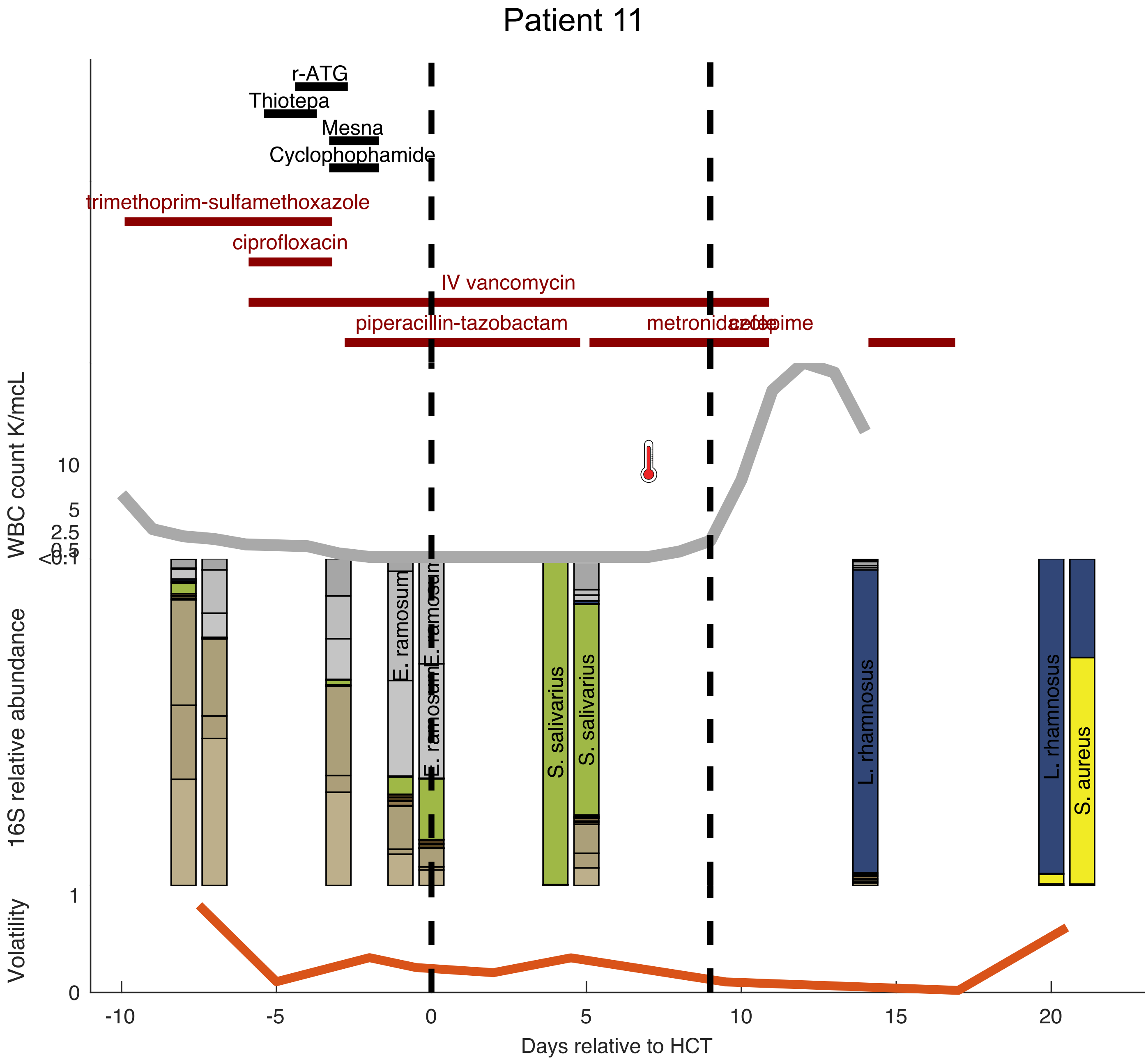

Patient 11
r-ATG
Thiotepa
Mesna
Cyclophophamide
trimethoprim-sulfamethoxazole
ciprofloxacin
IV vancomycin
piperacillin-tazobactam
metronidazole
cefepime
WBC count K/mcL
10
5
2.5
0.5
<0.1
16S relative abundance
E. ramosum
E. ramosum
S. salivarius
S. salivarius
L. rhamnosus
L. rhamnosus
S. aureus
Volatility
1
0
-10
-5
0
5
10
15
20
Days relative to HCT

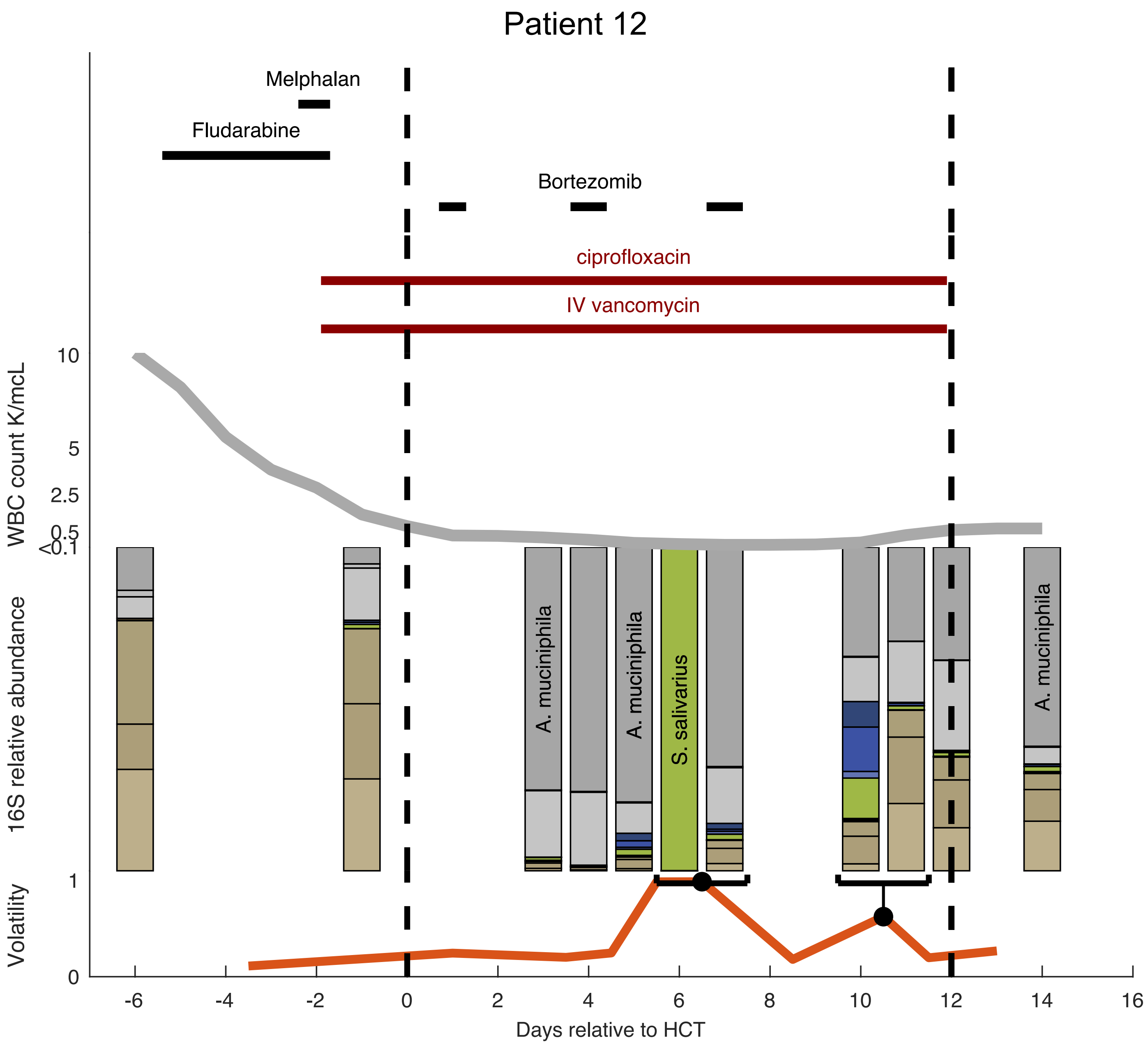

Patient 12
Melphalan
Fludarabine
Bortezomib
ciprofloxacin
IV vancomycin
WBC count K/mcL
10
5
2.5
0.5
<0.1
16S relative abundance
A. muciniphila
A. muciniphila
S. salivarius
A. muciniphila
Volatility
1
0
-6
-4
-2
0
2
4
6
8
10
12
14
16
Days relative to HCT

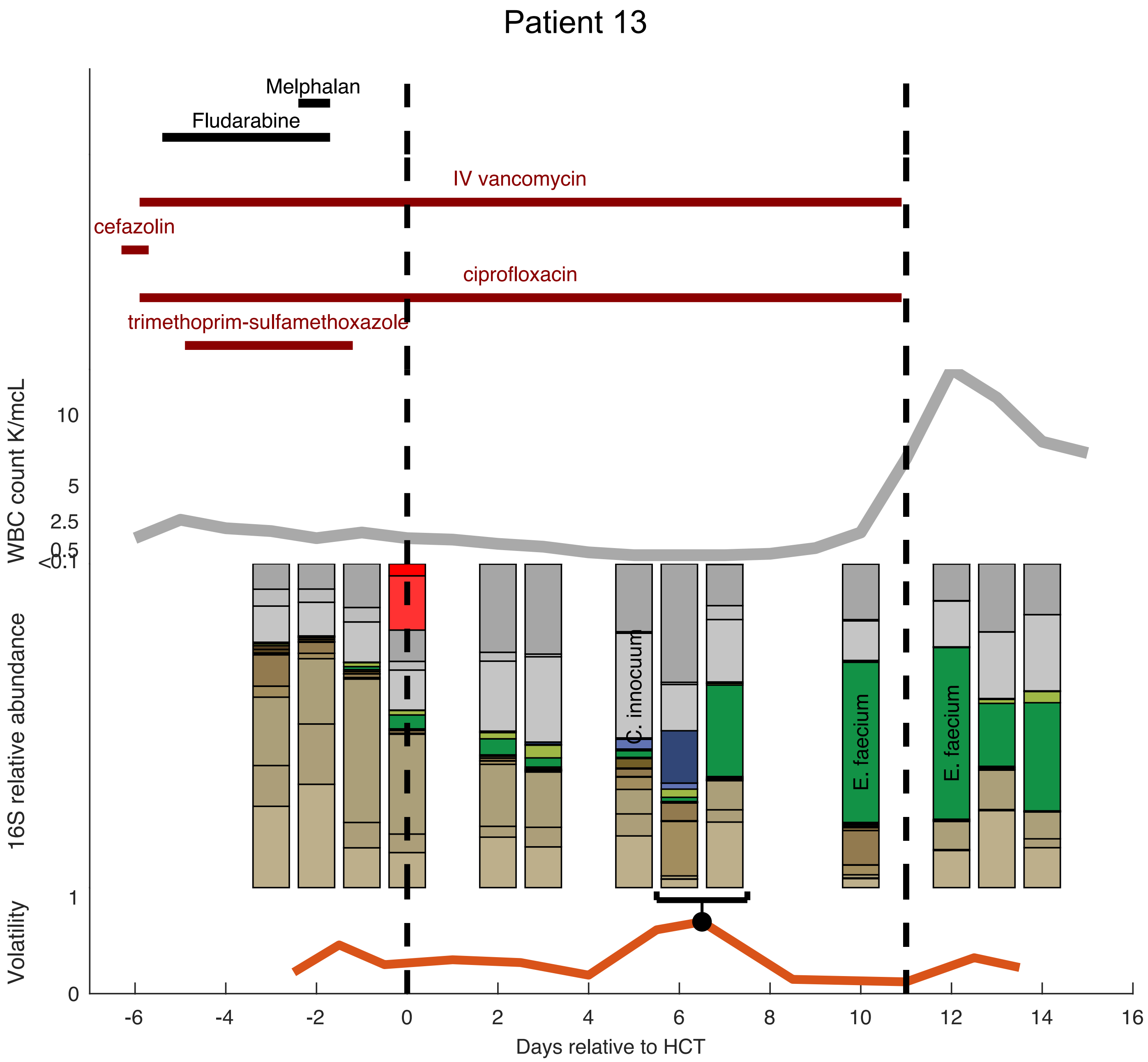

Patient 13
Melphalan
Fludarabine
IV vancomycin
cefazolin
ciprofloxacin
trimethoprim-sulfamethoxazole
WBC count K/mcL
10
5
2.5
0.5
<0.1
16S relative abundance
C. innocuum
E. faecium
E. faecium
Volatility
1
0
-6
-4
-2
0
2
4
6
8
10
12
14
16
Days relative to HCT

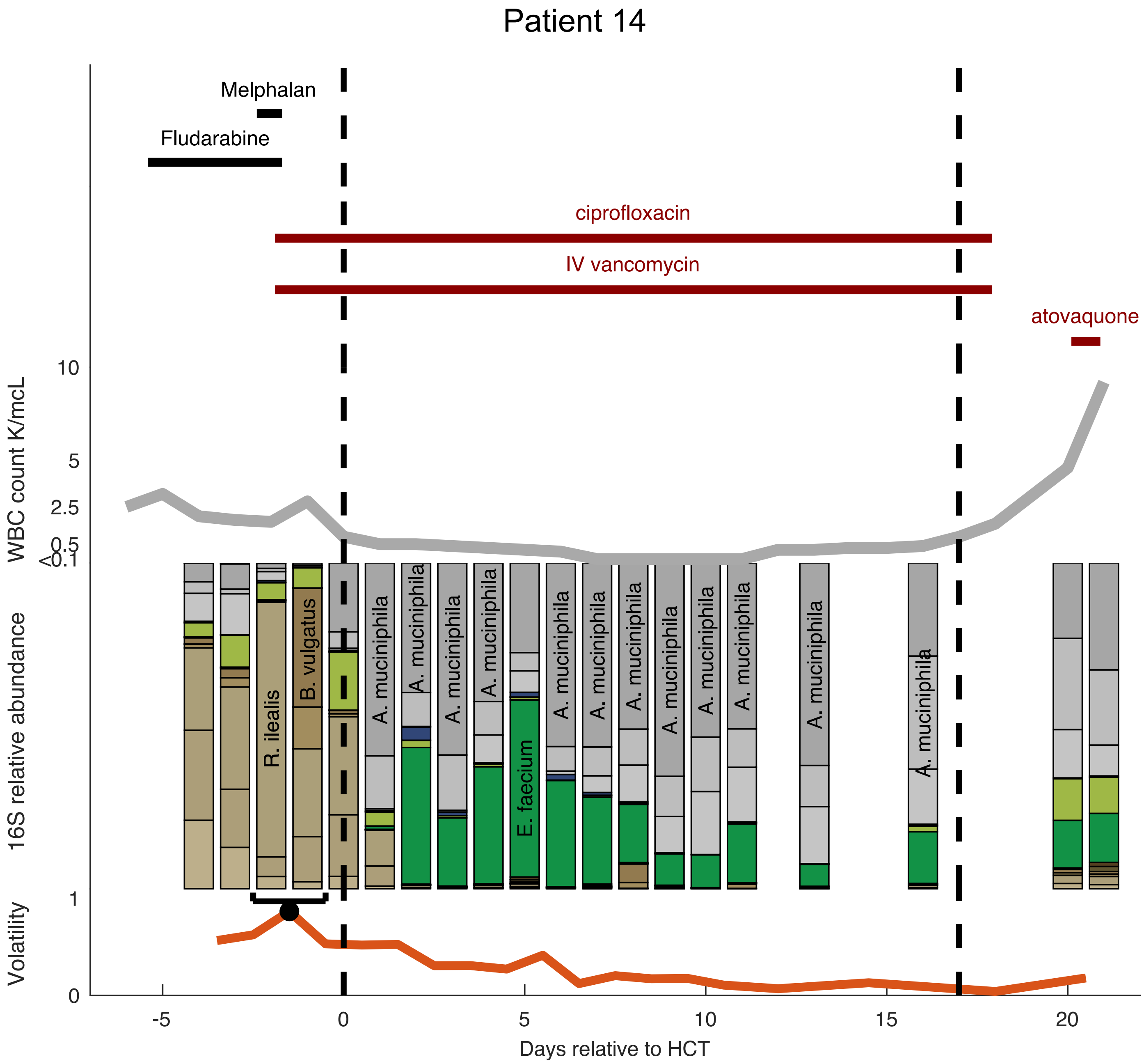

Patient 14
Melphalan
Fludarabine
ciprofloxacin
IV vancomycin
atovaquone
10
5
2.5
0.5
<0.1
WBC count K/mcL
16S relative abundance
R. ilealis
B. vulgatus
A. muciniphila
E. faecium
1
0
Volatility
-5
0
5
10
15
20
Days relative to HCT

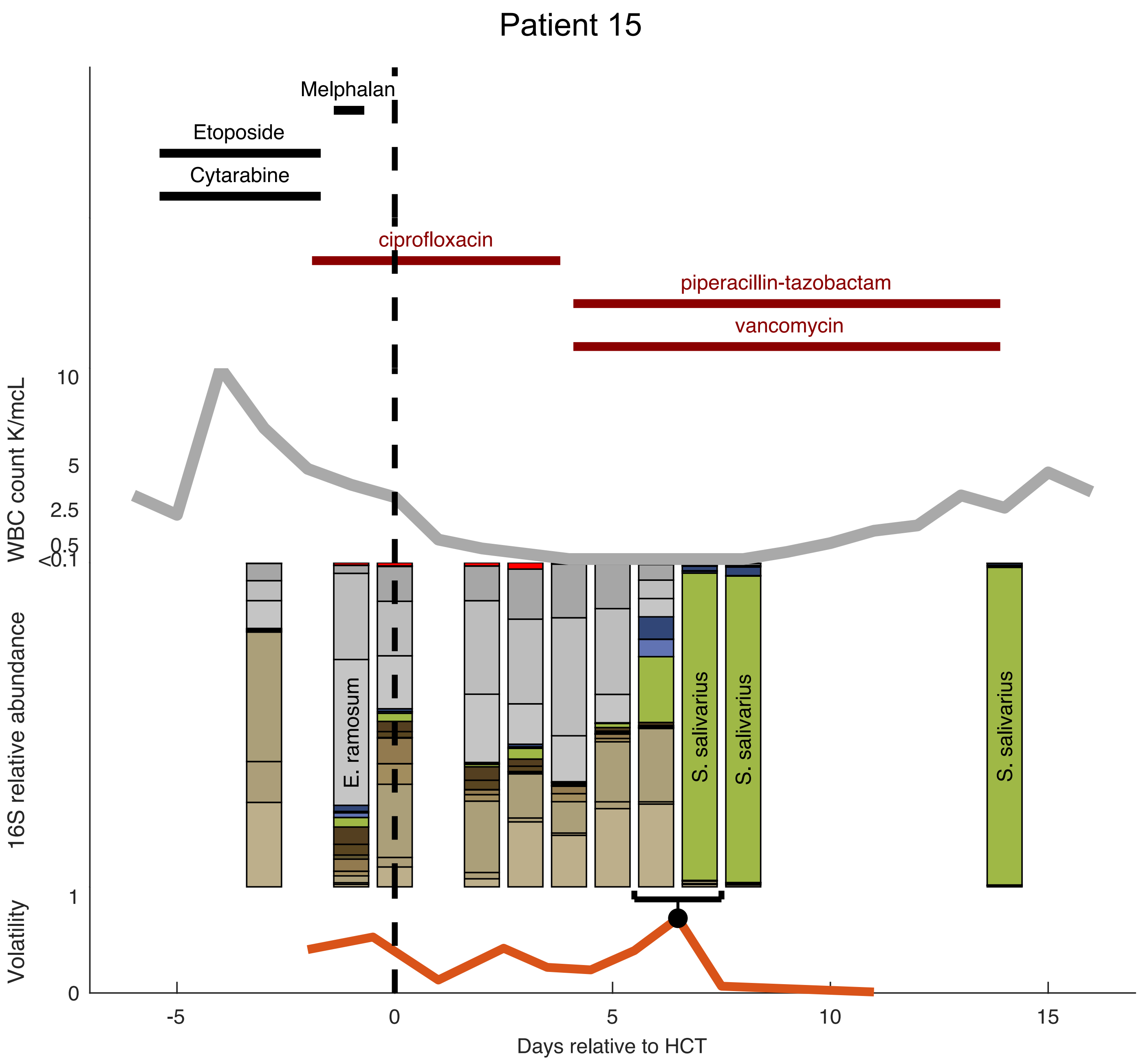
Patient 15
Melphalan
Etoposide
Cytarabine
ciprofloxacin
piperacillin-tazobactam
vancomycin
WBC count K/mcL
10
5
2.5
0.5
<0.1
16S relative abundance
E. ramosum
S. salivarius
S. salivarius
S. salivarius
Volatility
1
0
-5
0
5
10
15
Days relative to HCT

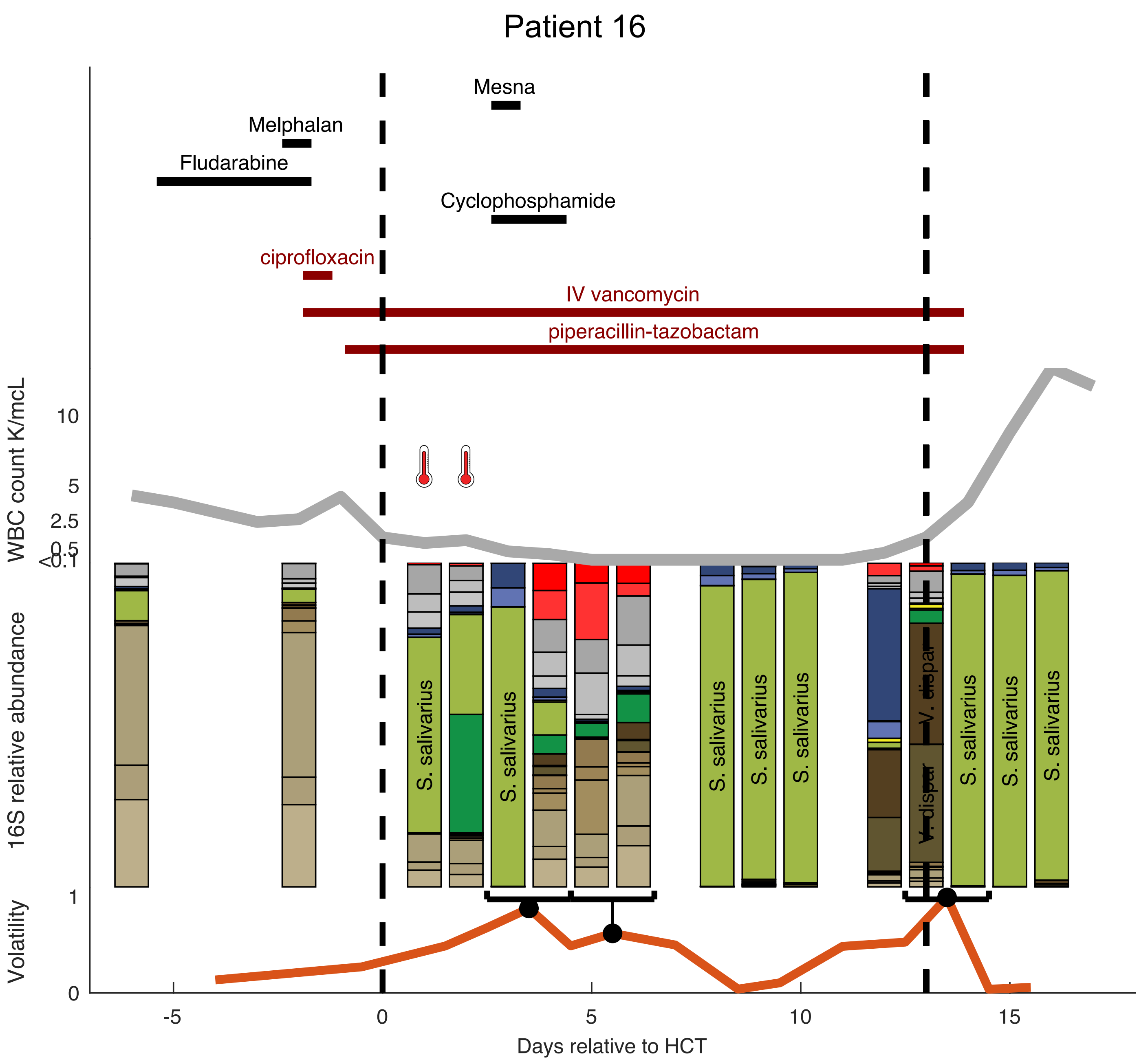

Patient 16
Mesna
Melphalan
Fludarabine
Cyclophosphamide
ciprofloxacin
IV vancomycin
piperacillin-tazobactam
WBC count K/mcL
10
5
2.5
0.5
<0.1
16S relative abundance
S. salivarius
V. dispar
Volatility
1
0
-5
0
5
10
15
Days relative to HCT

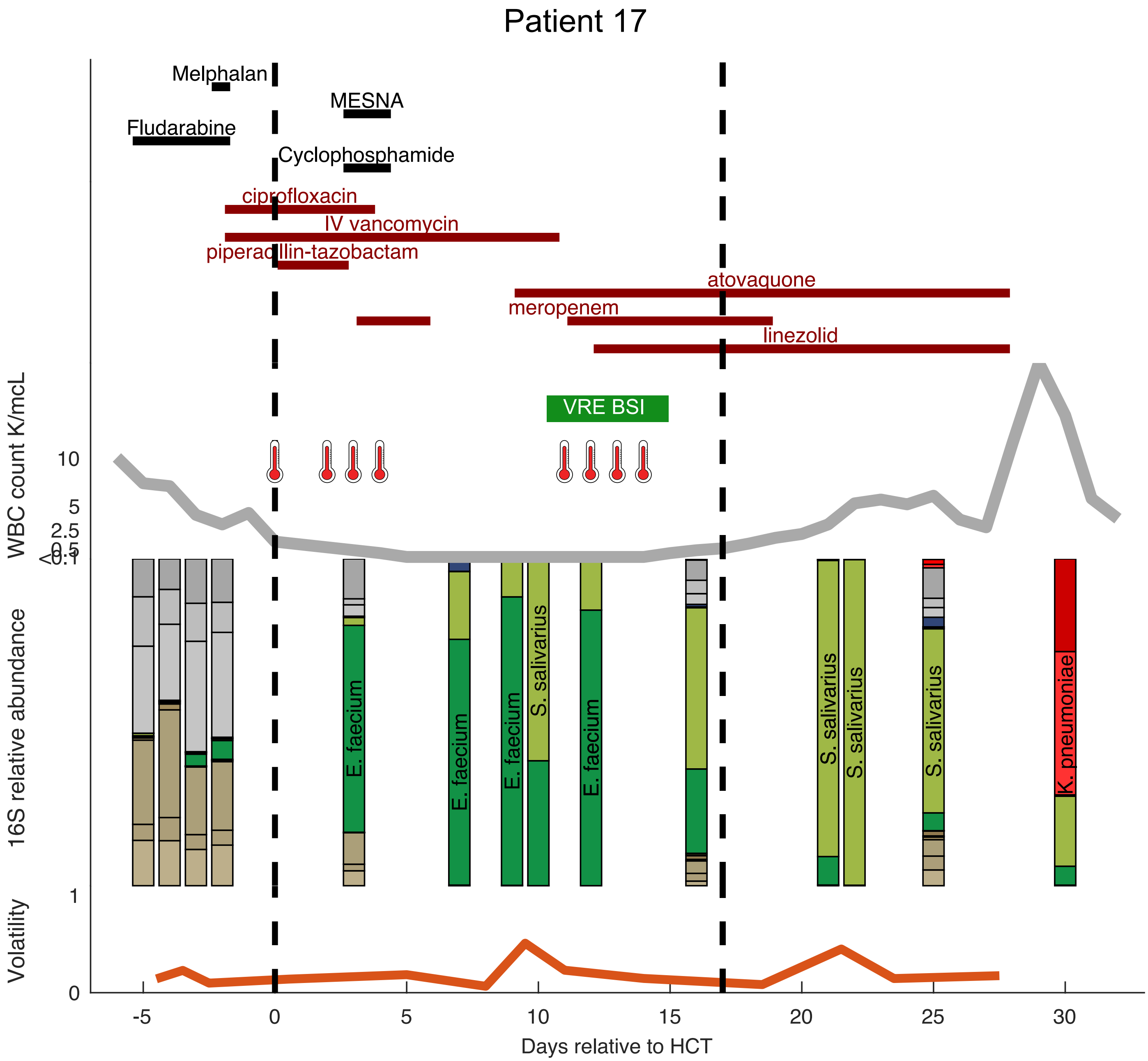
Patient 17
Melphalan
MESNA
Fludarabine
Cyclophosphamide
ciprofloxacin
IV vancomycin
piperacillin-tazobactam
atovaquone
meropenem
linezolid
VRE BSI
WBC count K/mcL
10
5
2.5
0.5
<0.1
16S relative abundance
E. faecium
E. faecium
E. faecium
S. salivarius
E. faecium
S. salivarius
S. salivarius
S. salivarius
K. pneumoniae
Volatility
1
0
-5
0
5
10
15
20
25
30
Days relative to HCT

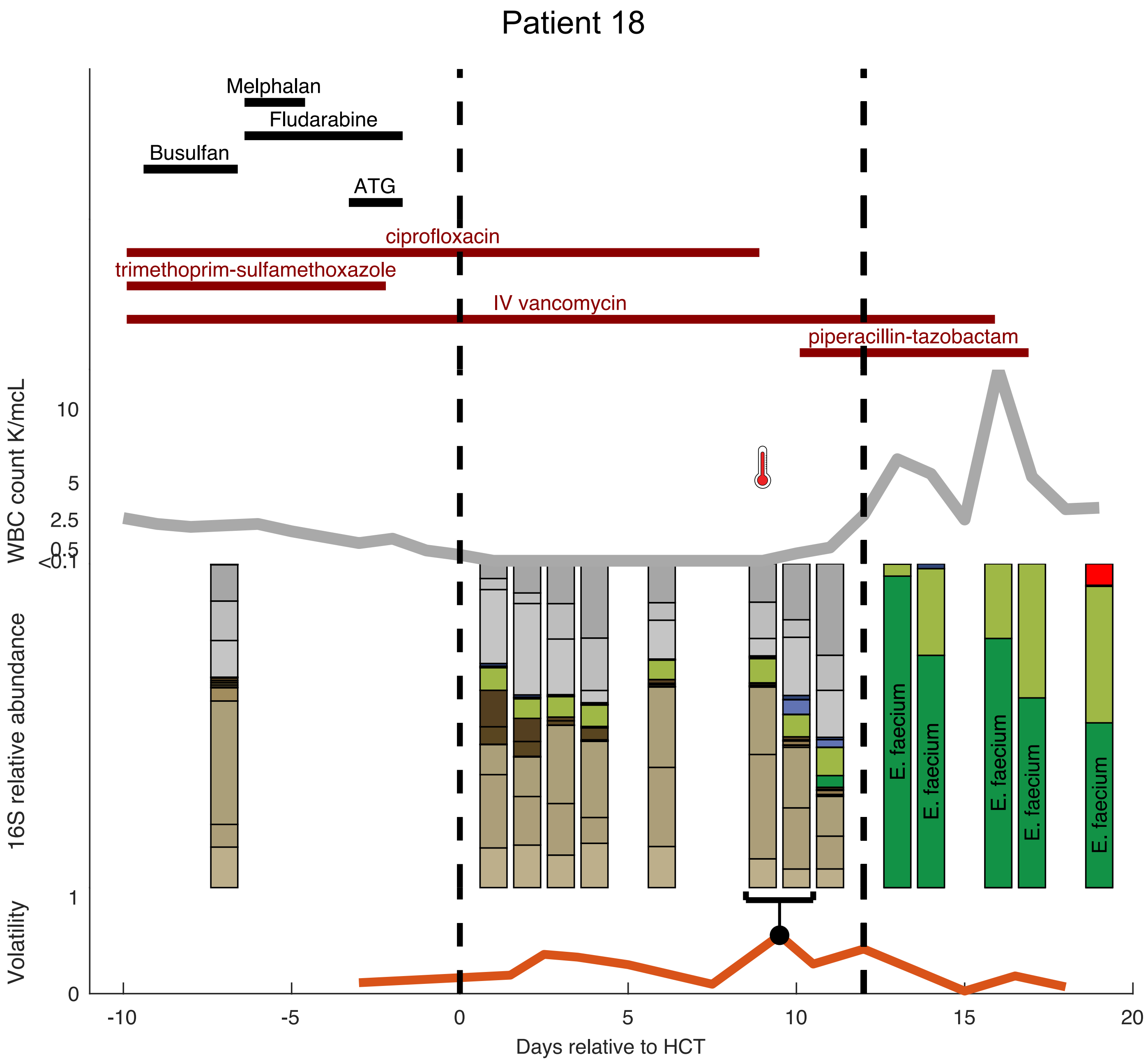
Patient 18
Melphalan
Fludarabine
Busulfan
ATG
ciprofloxacin
trimethoprim-sulfamethoxazole
IV vancomycin
piperacillin-tazobactam
WBC count K/mcL
10
5
2.5
0.5
<0.1
16S relative abundance
E. faecium
E. faecium
E. faecium
E. faecium
E. faecium
Volatility
1
0
-10
-5
0
5
10
15
20
Days relative to HCT
